# A Complete Set of Equations and Parameters for the Computational Model of Mitochondrial Function for the Proximal Convoluted Tubule and Medullary Thick Ascending Limb Cells in the Rat Kidney

**DOI:** 10.1101/2022.01.27.477845

**Authors:** T. William Bell, Anita T. Layton

## Abstract

To investigate mitochondrial function of renal epithelial cells under different tissue oxygenation levels, we have developed and applied computational models of mitochondrial function of the proximal convoluted tubule and medullary thick ascending limb cells. The models predict several key cellular quantities, including ATP generation, P/O (phosphate/oxygen) ratio, proton motive force, electrical potential gradient, oxygen consumption, the redox state of important electron carriers, and ATP consumption. The complete set of model equations and parameters are presented here, together with local and global sensitivity analysis results.

## 1 Model Equations

We present a computational model of mitochondrial function of renal epithelial cells. The model includes 53 state variables representing concentrations in three compartments, and the electrical potential gradient across the inner membrane of the mitochondrion. We include the full table of state variables (see Table 12), a table of adjustable parameters for the cytosol, membrane fluxes, and buffering reactions (see Table 13), and a table of physical parameters (see Table 14 at the end of this chapter, we characterize less important kinetic parameters in the section with the relevant flux or reaction in a table (for further discussion see the supplementary material to Wu et al. [10], our exposition draws heavily on Wu et al.). Our model includes the process of pyruvate oxidation, the TCA cycle, and oxidative phosphorylation. Aside from this we also include a couple significant reactions in the matrix and cytosol, several transporter activities on the inner membrane of the mitochondrion, and several passive diffusive fluxes across the outer membrane of the mitochondrion. We begin by showing our model from a bird’s eye view, with the fluxes and reactions between state variables represented but with the details of the kinetics left out. Then we will dive into the exact kinetics after that. In the work of Wu et al. [10], 35 parameters were estimated. Many more parameters are taken from direct experimental estimates as discussed in the supplementary material to Wu et al [10]. In our model, we include a new enzyme activity, glutamate dehydrogenase. For that enzyme we use experimental estimates for both its activity and Michaelis constant [8]. Since we are considering two novel tissues we identify experimental measurements for several key parameters that are specific to those tissues, and we fit or refit multiple parameters. Each chapter includes a table listing experimental sources for parameters that were changed based on direct measurements. Other parameter changes are discussed in the text itself.

**Table 1:** Kinetic parameters for pyruvate dehydrogenase.

| Parameter | Model Value ($\mu\text{M}$) |
| --- | --- |
| $K_{mA}$ | 38.3 |
| $K_{mB}$ | 9.9 |
| $K_{mC}$ | 60.7 |
| $K_{iNADH}$ | 40.2 |
| $K_{iACCOA}$ | 40.0 |

The electrical potential gradient depends on the difference in charges across the inner membrane of the mitochondrion, and drives the activity of ATP synthase. The gradient is determined by the net flux across the inner membrane of the mitochondrion, in our model the net charge across the inner membrane is influenced by the members of the electron transport chain (*J*_C1_, *J*_C3_, and *J*_C4_), hydrogen leak (*J*_Hle_), ATP synthase (*J*_F1_), and adenosine nucleotide translocase (*J*_ANT_). Notably, while the stoichiometry of all these fluxes are well-understood aside from ATP synthase, where the necessary hydrogen flux is controversial, thus the flux term for ATP synthase has a parameter *n_A_* that determines the stochiometry of the flux. In our model, *n_A_* is three. The electrical potential gradient is reported in millivolts.

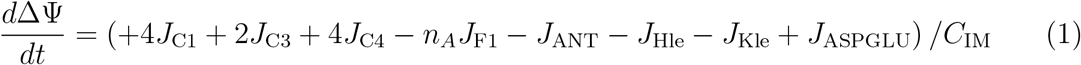

The proton motive force may be calculated using the electrical potential gradient and the hydrogen ion concentrations in the matrix and intermembrane space as follows:

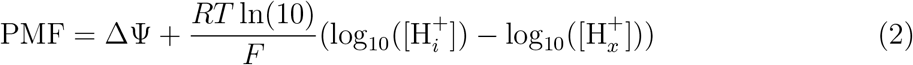

By far the most important compartment for our model is the mitochondrial matrix. The mitochondrial matrix includes all of the reactions involved in the TCA cycle (*J*_acon_, *J*_isod_, *J*_akgd_, *J*_scoas_, *J*_sdh_, *J*_fum_, *J*_mdh_, *J*_got_, and *J*_cits_) and pyruvate oxidation (*J*_pdh_). Aside from these reactions, there are several fluxes that we identify as significant, both the ones related to oxidative phosphorylation mentioned before, several transporters that control the quantities of TCA cycle intermediates within the mitochondrial matrix (*J*_PYRH_, *J*_MALPI_, *J*_SUCPI_, *J*_SUCMAL_, *J*_AKGMAL_, and *J*_CITMAL_ and several other transporters (*J*_PIHO_ *J*_ANT_ *J*_KH_ *J*_GLUH_ and *J*_ASPGLU_). The concentrations of certain essential compounds are fixed, most notably the oxygen concentration. We consider a range of oxygen concentrations in our work. In what follows, we use the subscript ‘x’ for matrix concentrations, ‘i’ for intermembrane space concentrations, and ‘c’ for cytosolic concentrations.

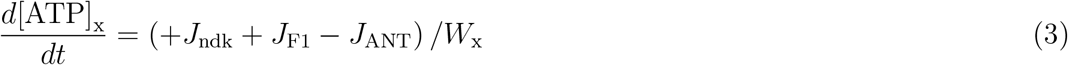

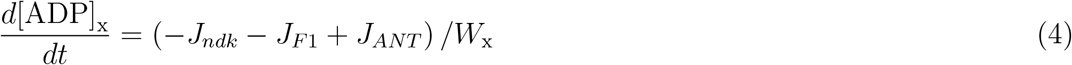

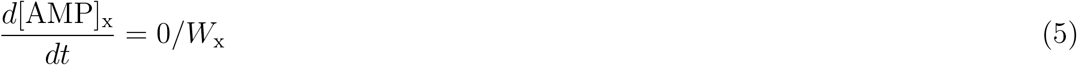

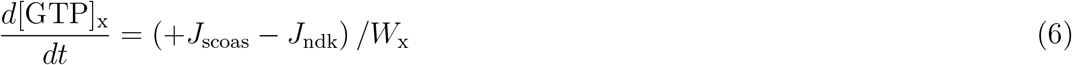

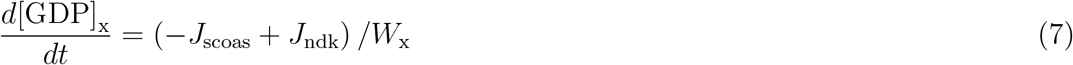

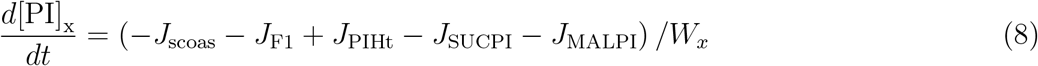

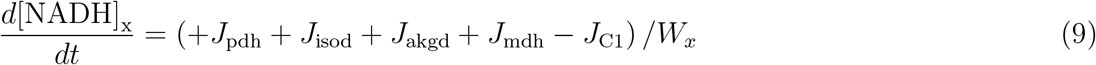

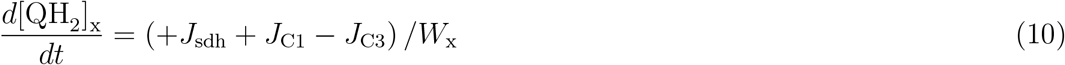

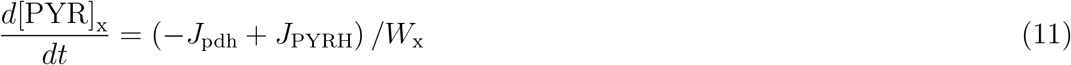

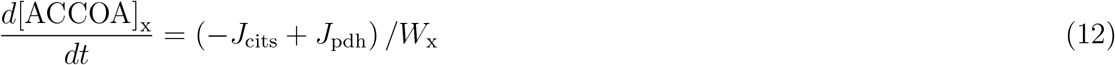

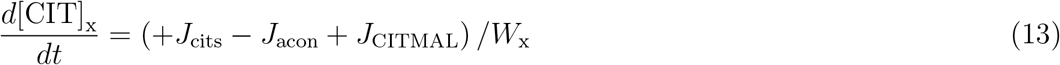

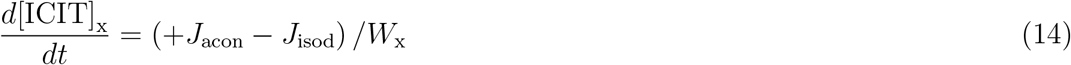

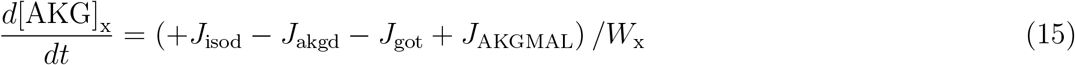

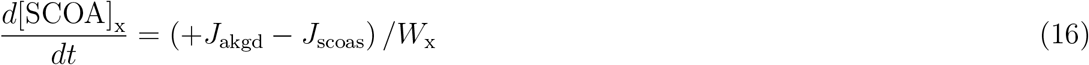

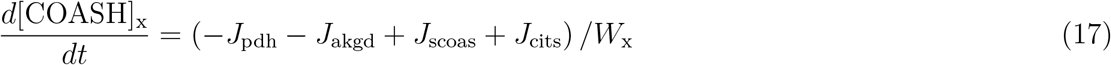

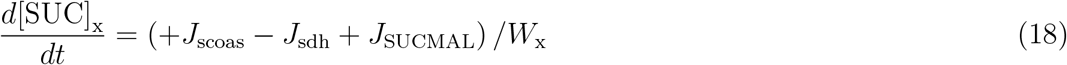

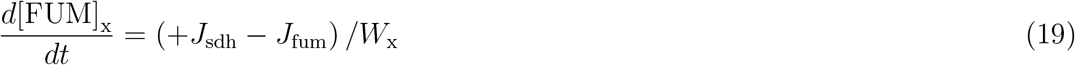

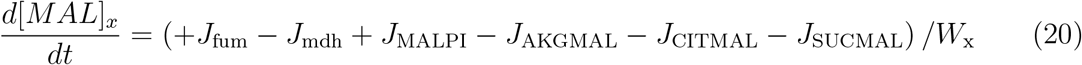

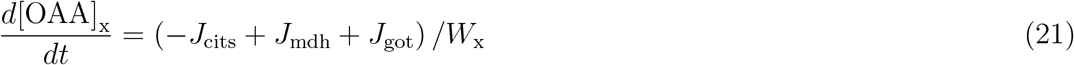

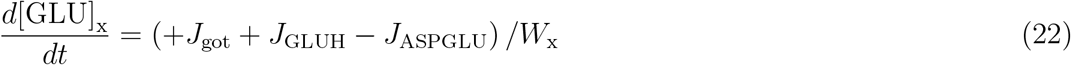

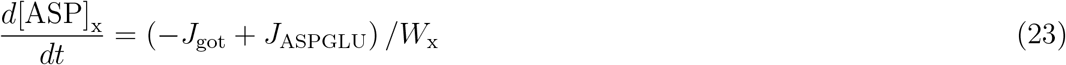

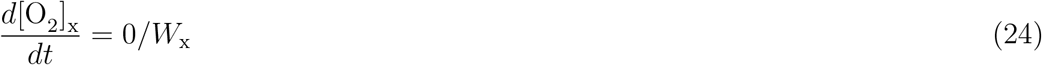

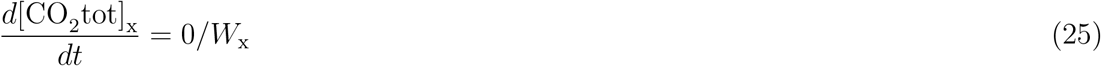

For the liver, our model includes the activity of glutamate dehydrogenase (*J*_gdh_), which converts glutamate into alpha-ketogluterate. With this adjustment the model has the following structure:

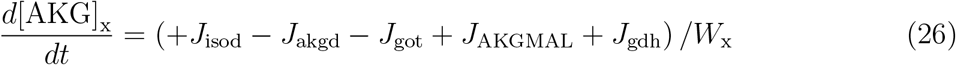

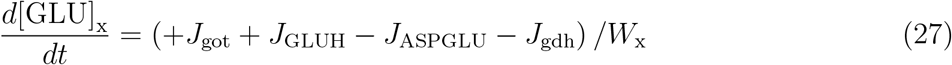

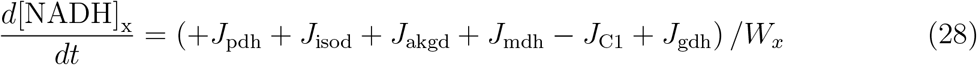

The intermembrane space connects the mitochondrial matrix and the cytosol (or for isolated mitochondria, the surrounding compartment). The inner membrane is relatively impermeable, relying on several transporters to maintain the flux of solutes between the matrix and the intermembrane space (transporters and oxidative phosphorylation complexes mentioned previously). On the other hand, the outer membrane is highly permeable, and so the intermembrane space & cytosol are coupled by passive diffusion in multiple solutes (*J*_ATP_, *J*_ADP_, *J*_AMPt_, *J*_PIt_, *J*_PYRt_, *J*_CITt_, *J*_ICITt_, *J*_AKGt_, *J*_SUCt_, *J*_MALt_, *J*_GLUt_, and *J*_ASPt_). These two connections give us the following equations:

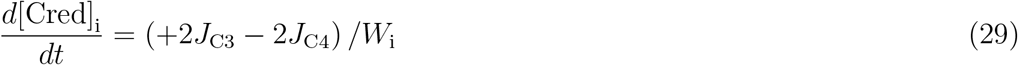

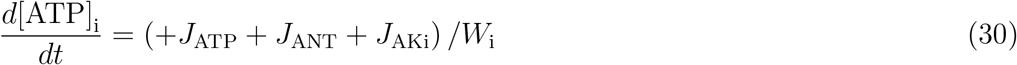

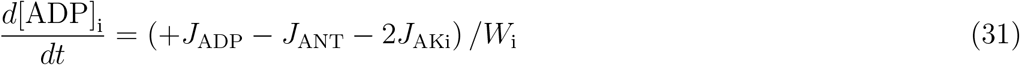

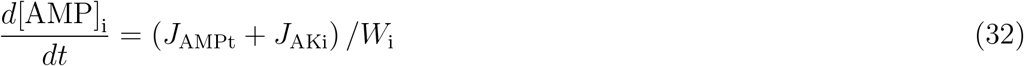

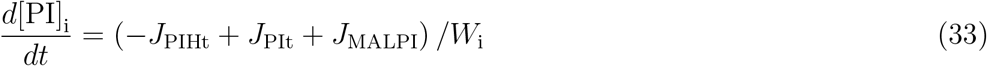

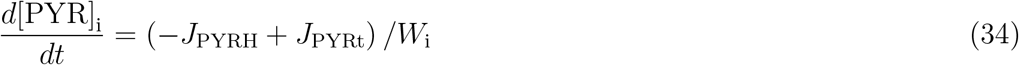

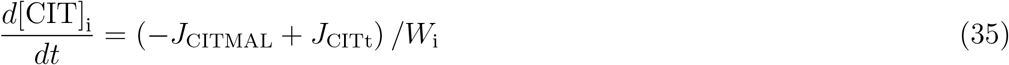

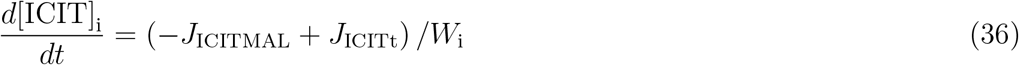

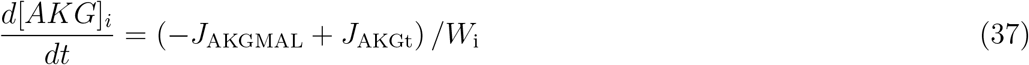

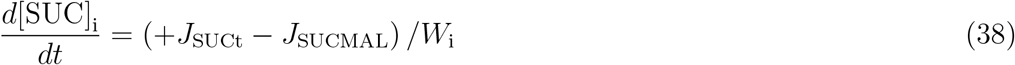

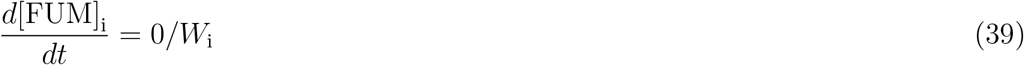

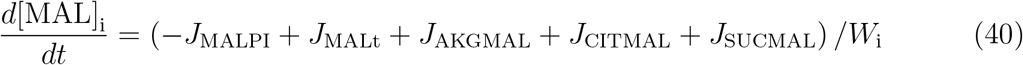

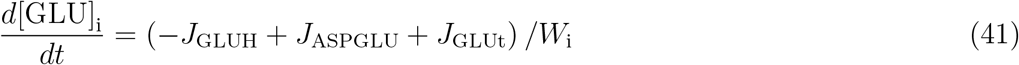

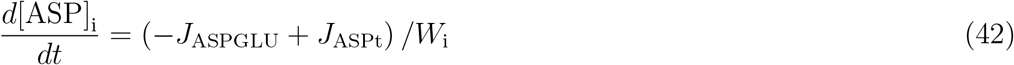

Our model covers two cases, in the first case, the outer compartment is the external space in which the isolated mitochondria are found. This case is important for several experimental frameworks used for measuring the P/O ratio, oxygen consumption, and hydrogen leak:

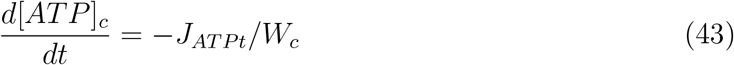

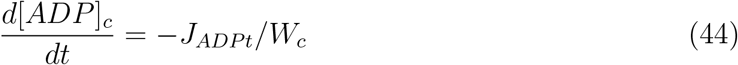

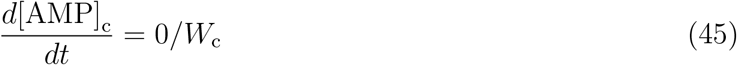

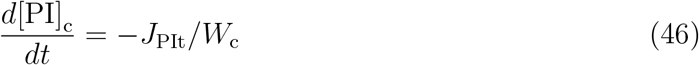

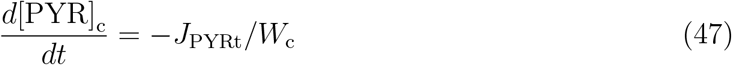

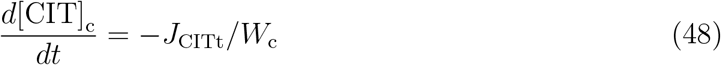

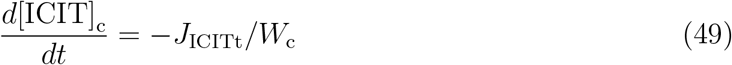

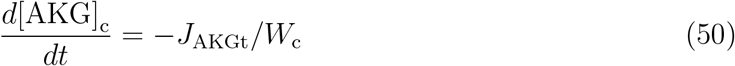

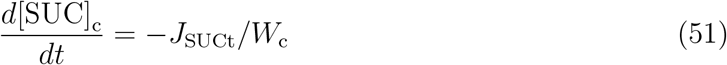

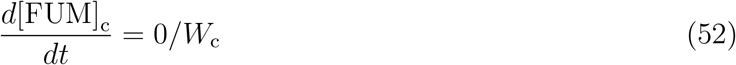

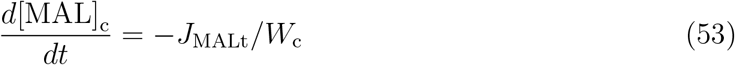

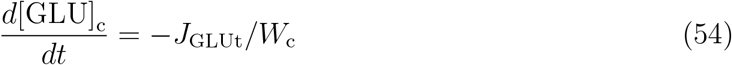

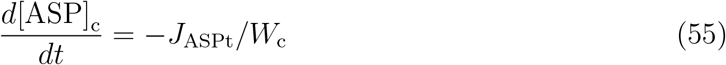

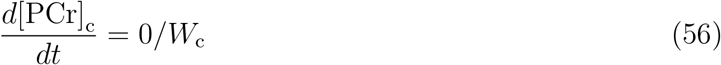

If the external space is the cytoplasm, then the system is forced to remain at its nonequilibrium steady state by clamping the concentration of pyruvate in the cytoplasm. This ensures that the mitochondrion will not exhaust its supply of pyruvate and that it will continue to produce ATP. Otherwise most compounds are determined by the diffusion across the outer mitochondrial membrane. The exceptions to this were the ATP consumption (*J*_AtC_), in addition, ADP may phosphorylate ADP, producing ATP and AMP (*J*_AKe_), and the creatine kinase reaction which consumes creatine and ADP to produce ATP and hydrogen ions (*J*_CKe_).

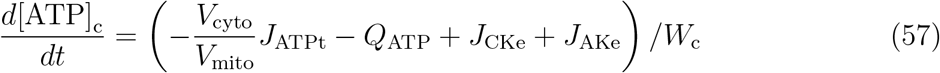

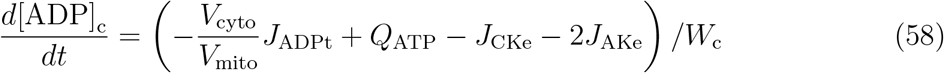

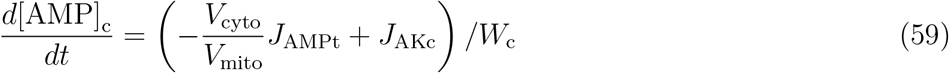

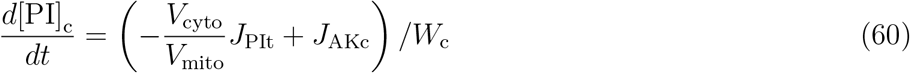

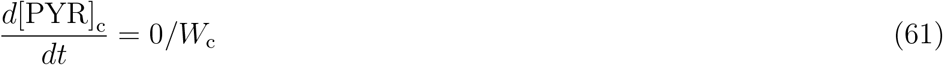

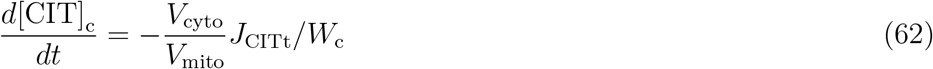

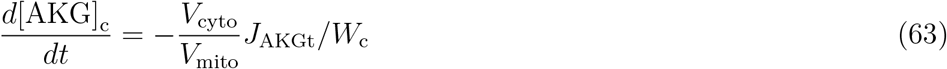

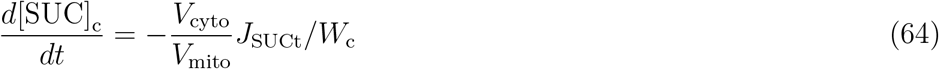

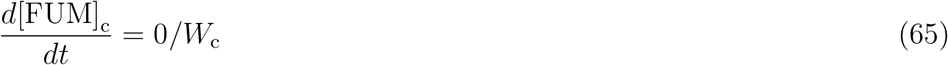

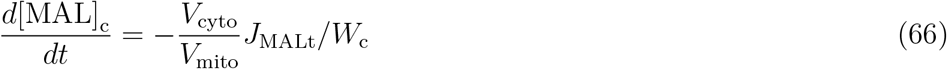

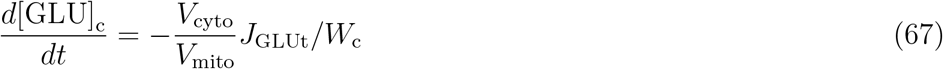

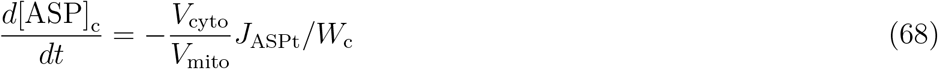

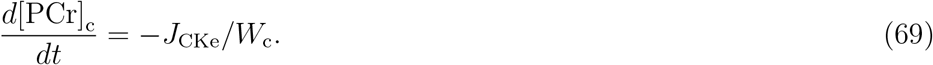

Some quantities were calculated explicitly from the above differential equations, assuming that the sum of two concentrations were conserved. This required us to find the pooled concentration of NAD^+^ and NADH, of ubiquinone (oxidized coenzyme Q) and ubiquinol (reduced coenzyme Q), and Cytochrome C in both its reduced & oxidized forms. This gives us the following additional equations:

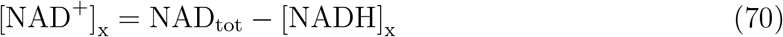

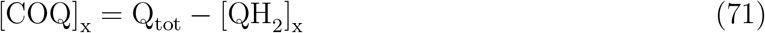

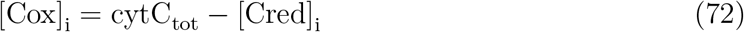

So far we have ignored the ion concentrations of hydrogen, magnesium, and potassium. These ion concentrations are fixed in the intermembrane space and cytosol. In the mitochondrial matrix these quantities are not fixed. Their kinetics are complex because there are binding sites that the ions compete for and the mitochondrial matrix is buffered. Here we provide the equations describing their kinetics, and several quantities to simplify the equations. This model is derived in Beard and Qian [1] in the chapter “Biochemical Reaction Networks”. Starting with the big picture, we have the following expressions for the derivatives of the ion concentrations, whose terms we will write out explicitly below:

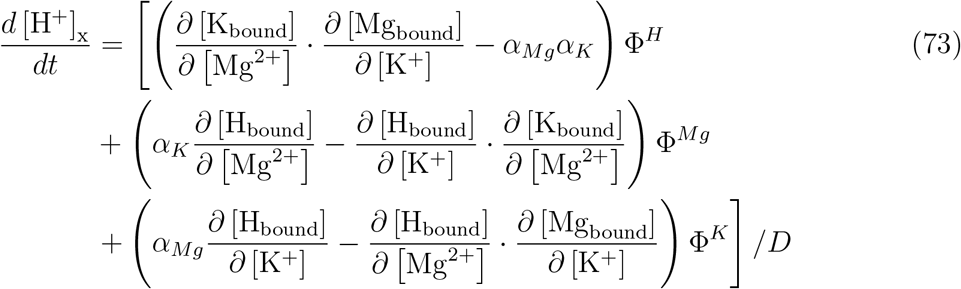

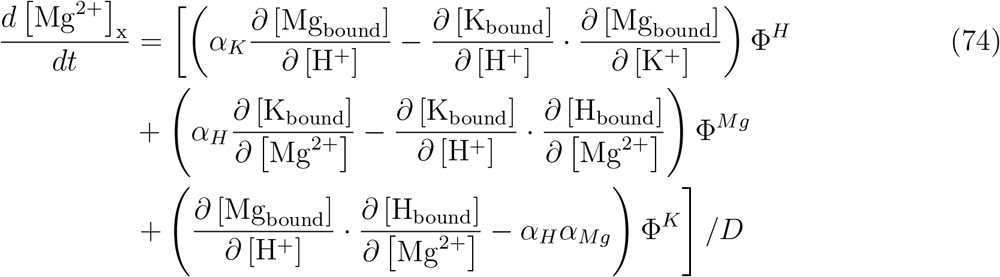

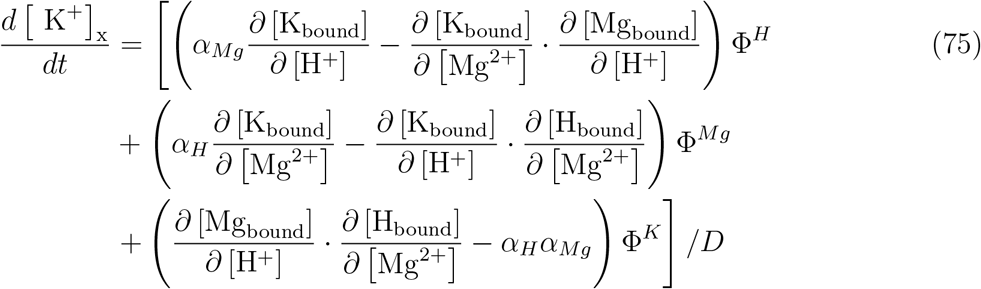

We’ll start by unpacking the expressions that are not partial derivatives. These include the denominator, the expressions connecting the binding to the cross-membrane fluxes of the model (Φ^*H*^, Φ^*Mg*^, and Φ^*K*^), and the buffer terms (*α*_*H*_, *α_Mg_*, and *α_K_*). In what follows, L; under a summation refers to a generic reactant with hydrogen or the two metal ions considered, the full list of such reactants includes ATP, ADP, AMP, GTP, phosphate, NADH/NAD^+^, coenzyme Q, oxaloacetate, acetyl-CoA, citrate, isocitrate, alphaketogluterate, succinyl-CoA, coenzyme A, succinate, fumerate, malate, glutamate, aspartate, cytochrome C, oxygen, water, FADH_2_/FAD^2+^, carbon dioxide, phosphocreatinine, creatinine, pyruvate, glucose, and glucose-6-phosphate. These reactants will also appear in the partial derivative expressions below.

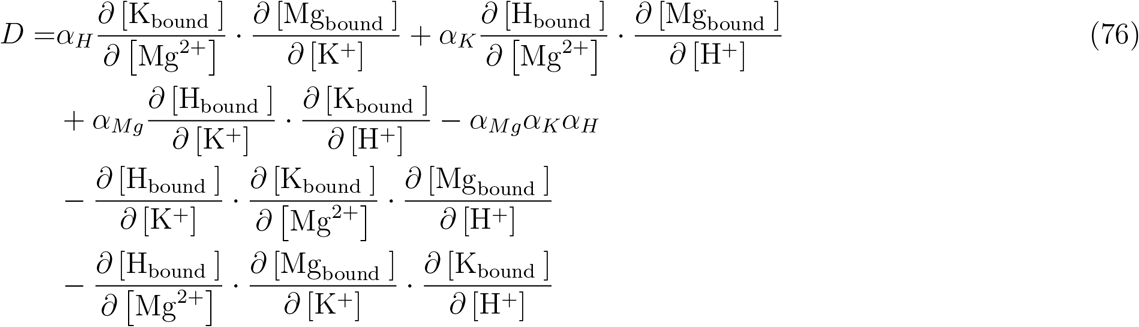

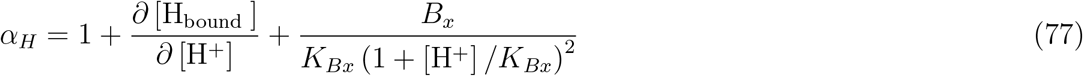

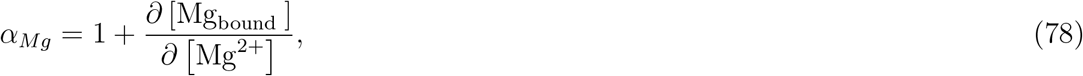

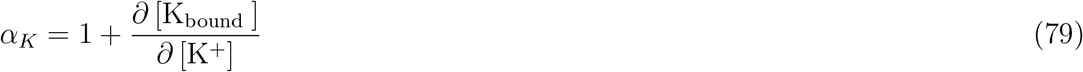

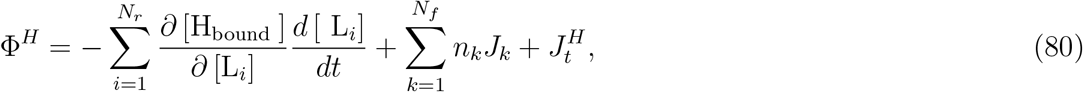

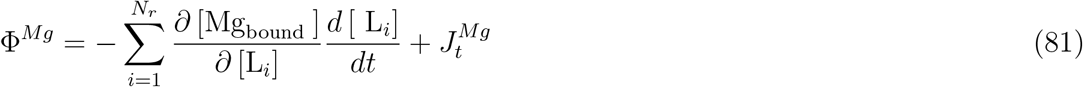

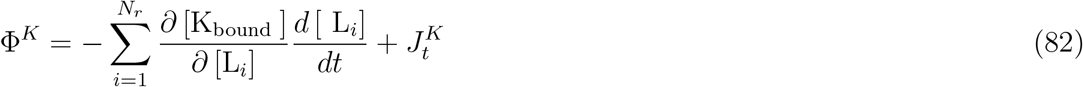

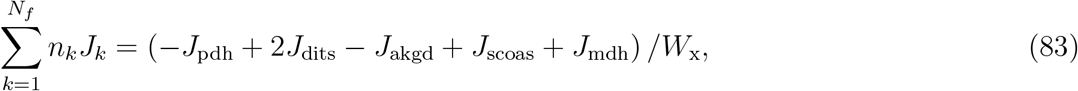

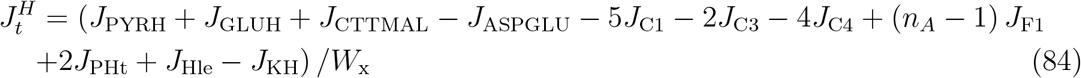

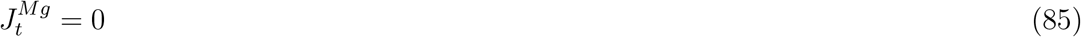

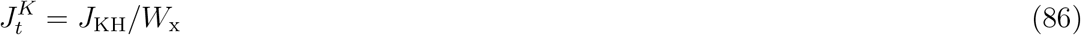

The remaining terms to define are partial derivatives of bound ions relative to the free concentration of each of the other above ions. These establish the extent to which an increase in the concentration of one ion will impact the binding opportunities for another ion.

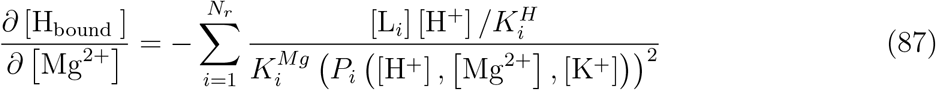

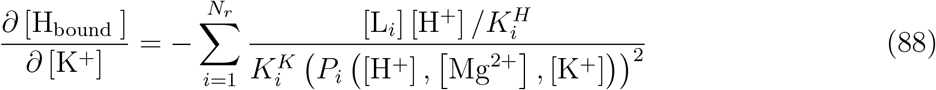

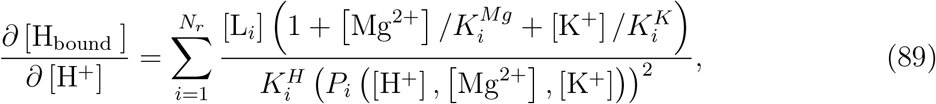

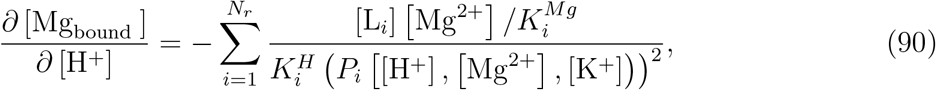

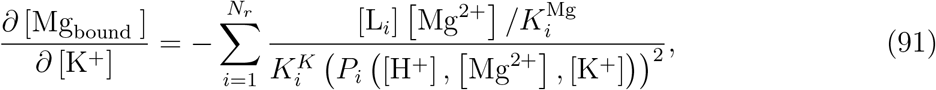

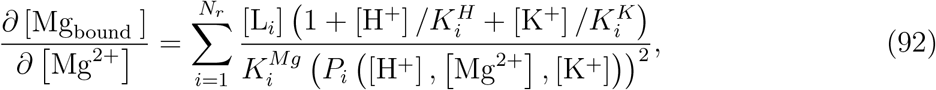

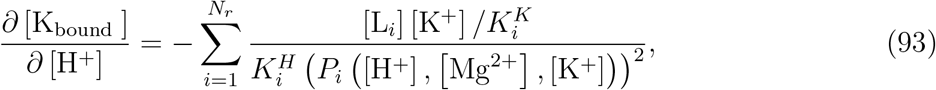

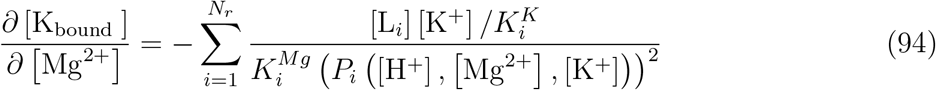

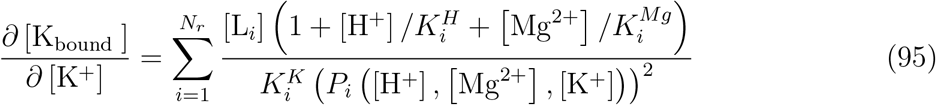

where *P_i_*, ([H^+^], [Mg^2+^], [K^+^]) is the reaction polynomial which may be expressed as follows:

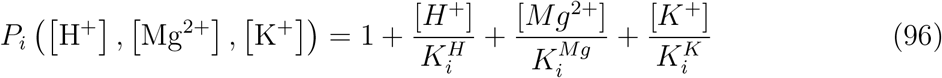

The complexity of ion binding is due to the many charged particles in the mitochondrial matrix to which these metal ions and protons may bind, and the complex buffering of ions in the cell.

### 1.1 Matrix Reactions

In this section we deal with reactions happening in the mitochondrial matrix, the majority of these reactions are directly related to the TCA cycle or pyruvate oxidation. Both biochemical pathways are subject to significant regulatory complexities, we will point it out where it occurs and where it is important we will indicate where the kinetics of the enzymes involved are discussed more at-length.

Pyruvate in the mitochondrial matrix is consumed by pyruvate dehydrogenase alongside coenzyme A and NAD^+^ to produce acetyl-CoA, NADH, and carbon dioxide. Pyruvate dehydrogenase may be phosphorylated by pyruvate dehydrogenase kinase, inhibiting pyruvate dehydrogenase. Pyruvate dehydrogenase kinase itself is activated by NADH and acetyl-CoA, and thus pyruvate dehydrogenase activity is inhibited by both. The kinetics of pyruvate dehydrogenase are as follows:

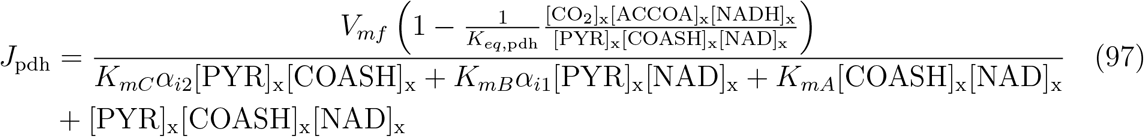

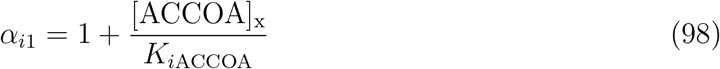

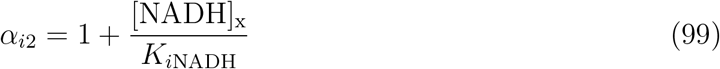

Pyruvate dehydrogenase kinetics are described at length in Cox & Nelson [3], and the reaction mechanism used to arrive at these kinetics are found in the appendix of Wu et al [10]. The parameters for *J*_pdh_ are summarized in Table 1. The form of the equilibrium constant here and below are discussed in the supplementary material for Wu et al. [10].

Acetyl-CoA produced by pyruvate dehydrogenase is consumed by citrate synthase along with oxaloacetate to produce citrate. Citrate synthase is also subject to complex regulation, including competitive inhibition by unchelated citrate of the oxaloacetate binding site, and uncompetitive inhibition by unchelated adenine nucleotides, unchelated coenzyme A, unchelated acetyl-CoA, and unchelated succinyl-CoA against the enzyme-acetyl-CoA complex, this gives us:

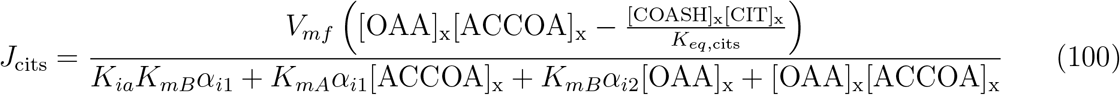

where *α*_*i*1_ and *α*_*i*2_ are as follows:

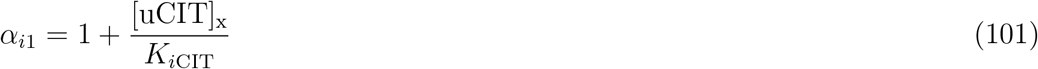

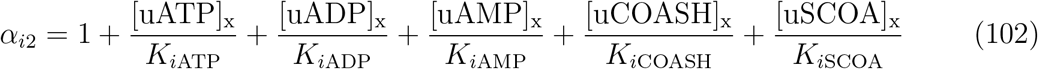

Here the prefix indicates that the compounds are unchelated. The mechanism underlying this model of citrate synthase is a mechanism described by Wu et al [10]. We give a list of parameters in Table 2.

**Table 2:** Kinetic parameters for citrate synthase.

| Parameter | Model Value ($\mu\text{M}$) |
| --- | --- |
| $K_{mA}$ | 4 |
| $K_{mB}$ | 14 |
| $K_{ia}$ | 3.33 |
| $K_{iCIT}$ | 1600 |
| $K_{iATP}$ | 900 |
| $K_{iADP}$ | 1800 |
| $K_{iAMP}$ | 6000 |
| $K_{iCOASH}$ | 67 |
| $K_{iSCOA}$ | 140 |

Acontinase converts citrate to isocitrate, and has a simple reaction mechanism. Naturally, its kinetics reflect this simplicity. It may be expressed as follows:

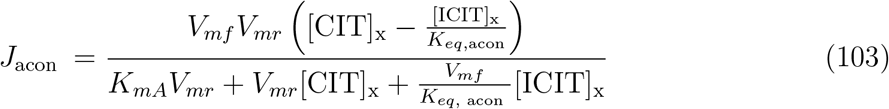

The parameters are catalogued in Table 3. *V_mr_* may be calculated as 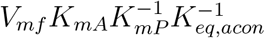.

**Table 3:**
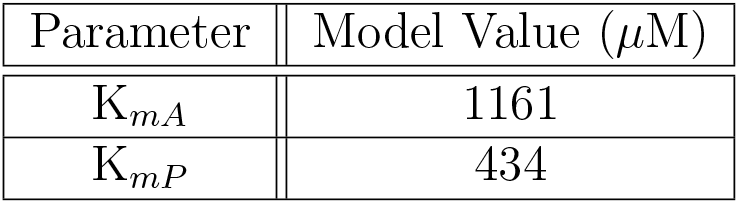
Kinetic parameters for acontinase.

| Parameter | Model Value ($\mu\text{M}$) |
| --- | --- |
| $K_{mA}$ | 1161 |
| $K_{mP}$ | 434 |

Isocitrate is then converted by isocitrate dehydrogenase to alpha-ketogluterate. In the process NAD^+^ is reduced to NADH, and carbon dioxide is produced. Free ATP acts as an allosteric inhibitor and free ADP may bind to the same site to prevent ATP-dependent inhibition (below we denote free ATP/ADP with a prefix f). NADH also competes for the binding site of NAD^+^. The reaction mechanism is discussed in the appendix of Wu et al. [10], and the regulation of the reaction is discussed at length in Kohn and Garfinkel [9]. Altogether we get the following expression for the flux via isocitrate dehydrogenase:

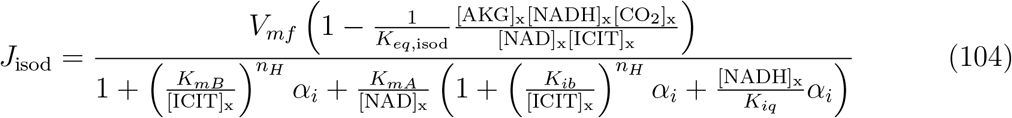

For the unitless exponent *n_H_*, we use the value 3 like in Wu et al [10]. Where the regulatory term *α_i_*, may be expressed as follows:

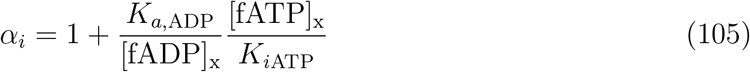

The kinetic parameters included are listed in Table 4.

**Table 4:**
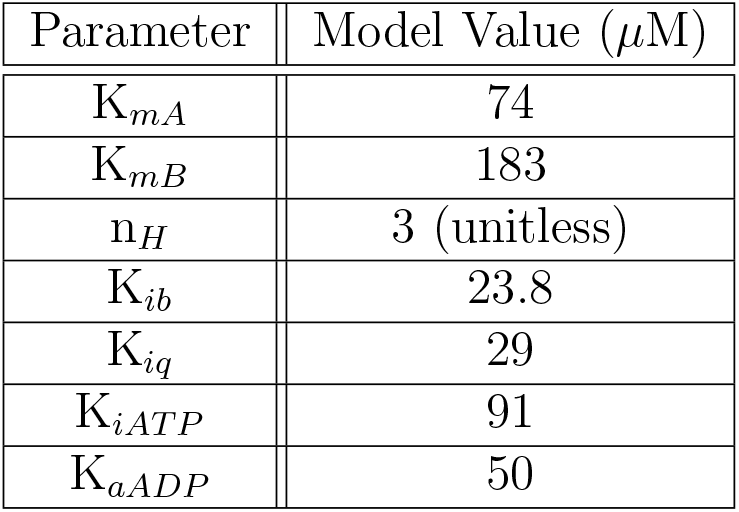
Kinetic parameters for isocitrate dehydrogenase.

| Parameter | Model Value ($\mu\text{M}$) |
| --- | --- |
| $K_{mA}$ | 74 |
| $K_{mB}$ | 183 |
| $n_H$ | 3 (unitless) |
| $K_{ib}$ | 23.8 |
| $K_{iq}$ | 29 |
| $K_{iATP}$ | 91 |
| $K_{aADP}$ | 50 |

Alpha-ketogluterate is used and produced by multiple reactions including the above reaction, in our exposition we will first introduce how it may be consumed to continue the TCA cycle. However we will return to alpha-ketogluterate again. Alpha-ketogluterate dehydrogenase consumes alpha-ketogluterate and continues the progression of intermediates through the TCA cycle. The reaction mechanism is discussed in the appendix of Wu et al. [10], and the inhibition-activation by adenine nucleotides is discussed by Kohn and Garfinkel [9]. The kinetics may be expressed as follows:

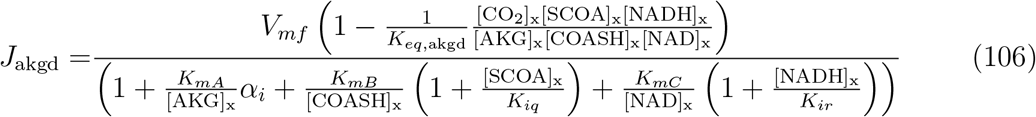

Where the regulatory term *α*_1_ may be expressed as follows:

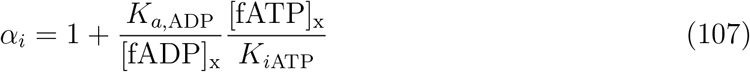

The kinetic parameters included are listed in Table 5.

**Table 5:** Kinetic parameters for alphaketogluterate dehydrogenase.

| Parameter | Model Value ($\mu\text{M}$) |
| --- | --- |
| $K_{mA}$ | 80 |
| $K_{mB}$ | 55 |
| $K_{mC}$ | 21 |
| $K_{iq}$ | 6.9 |
| $K_{ir}$ | 0.6 |
| $K_{iATP}$ | 50 |
| $K_{aADP}$ | 100 |

Alpha-ketogluterate is oxidized by the above reaction to succinyl-CoA, and in the process NAD^+^ is reduced to NADH. Succinyl-CoA is consumed by succinyl-CoA synthetase to produce succinate, in the process GDP is phosphorylated to GTP. Coenzyme A is also produced as a side product. While the reaction is not subject to serious regulation, the reaction catalysed by succinyl-CoA synthetase is quite complex and the expression for succinyl-CoA synthetase activity is as follows:

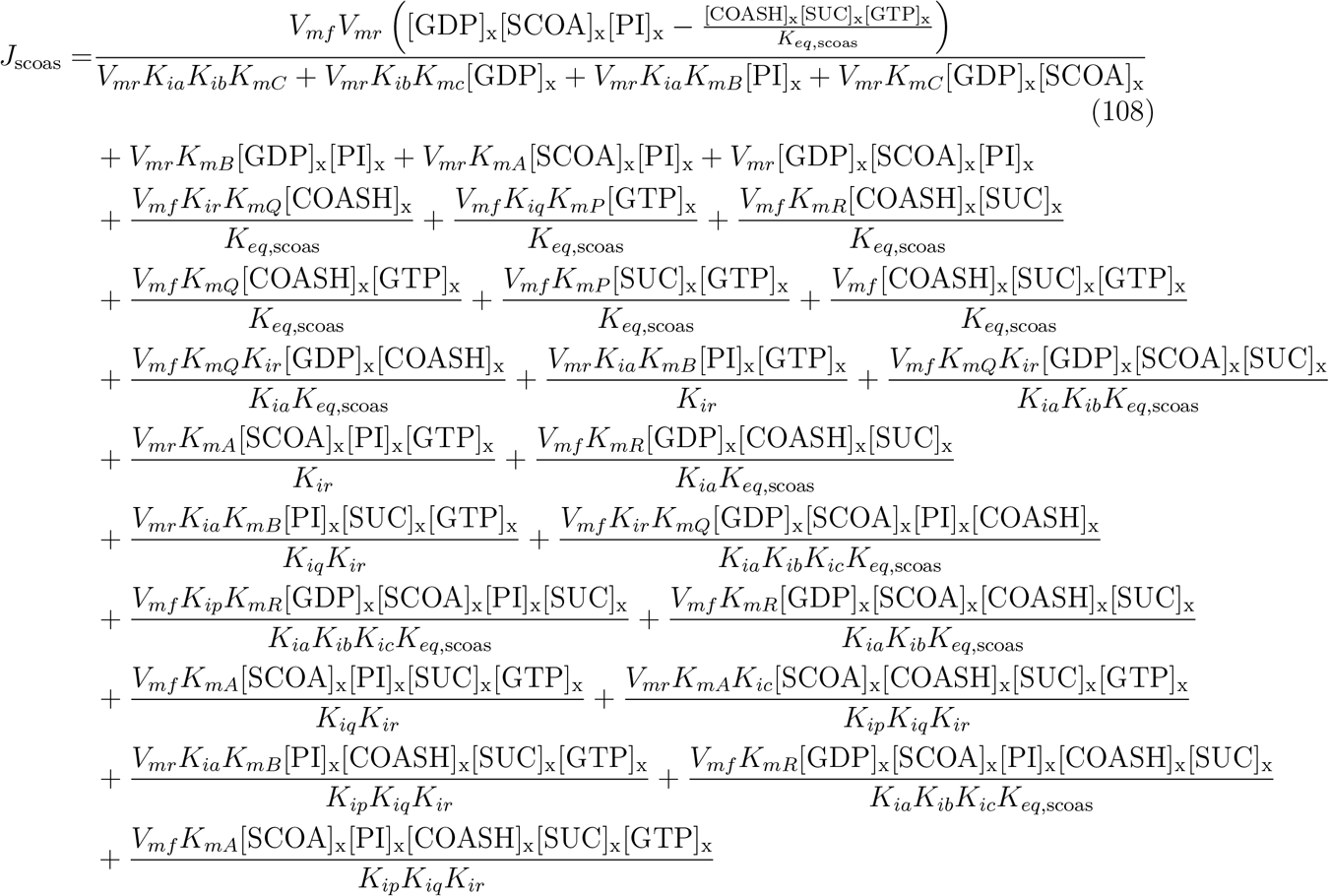

The parameters for this reaction are included in Table: The kinetic parameters included are listed in Table 6. *V_mr_* may be calculated as 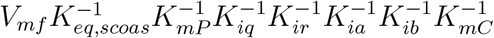.

**Table 6:** Kinetic parameters for succinyl-CoA synthetase.

| Parameter | Model Value ($\mu\text{M}$) |
| --- | --- |
| $K_{mA}$ | 16 |
| $K_{mB}$ | 55 |
| $K_{mC}$ | 660 |
| $K_{mP}$ | 20 |
| $K_{mQ}$ | 880 |
| $K_{mR}$ | 11.1 |
| $K_{ia}$ | 5.5 |
| $K_{ib}$ | 100 |
| $K_{ic}$ | 2000 |
| $K_{ip}$ | 20 |
| $K_{iq}$ | 3000 |
| $K_{ir}$ | 11.1 |

Succinate dehydrogenase, as well as being an enzyme involved in oxidative phosphorylation, directly catalyzes a step of the TCA cycle. Succinate from the above reaction is consumed by Complex II, allowing Complex II to donate an electron to coenzyme Q, an intermediate in oxidative phosphorylation, and producing fumarate in the process. The reaction mechanism assumed by the Kohn-Garfinkel model and used in Wu et al. [10] doesn’t explicitly include the implicit reduction and then oxidation of FAD/FADH_2_ that occurs in this process. The reaction is represented with the following kinetics:

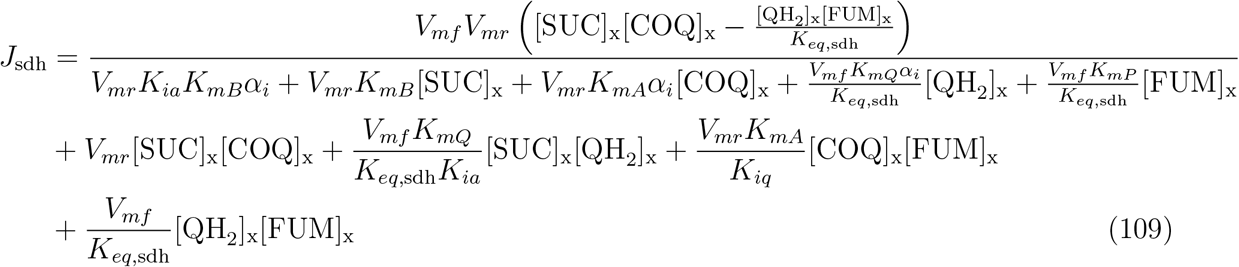

Where *α_i_*, is an inhibition-activation term, with activation by succinate and fumarate, and competitive inhibition of that activation by oxaloacetate. Altogether that gives us the following regulatory term *α_i_*:

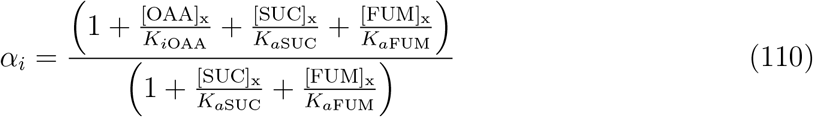

The parameters governing succinate dehydrogenase are found in Table 7. *V_mr_* can be calculated as 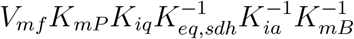

**Table 7:** Kinetic parameters for succinate dehydrogenase.

| Parameter | Model Value ($\mu\text{M}$) |
| --- | --- |
| $K_{mA}$ | 467 |
| $K_{mB}$ | 480 |
| $K_{mP}$ | 2.45 |
| $K_{mQ}$ | 1200 |
| $K_{ia}$ | 120 |
| $K_{iq}$ | 1275 |
| $K_{iOAA}$ | 1.5 |
| $K_{aSUC}$ | 450 |
| $K_{aFUM}$ | 375 |

Fumarate is in-turn consumed by fumarase to produce malate. Fumarase also binds to citrate, free adenine nucleotides, GTP, and GDP competitively. The mechanism is discussed in Wu et al. [10]:

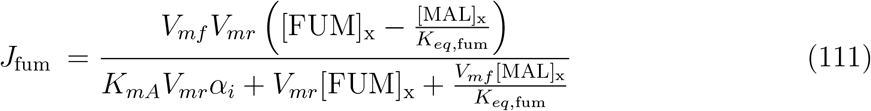

Where *α_i_* may be written as follows (as previously, the prefix f denotes that it is free, dissociated nucleotides):

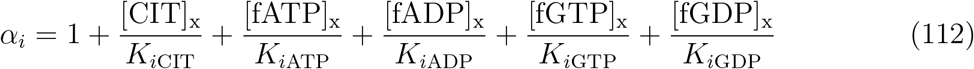

The kinetic parameters included are listed in Table 8. *V_mr_* is calculated as 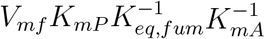.

**Table 8:**
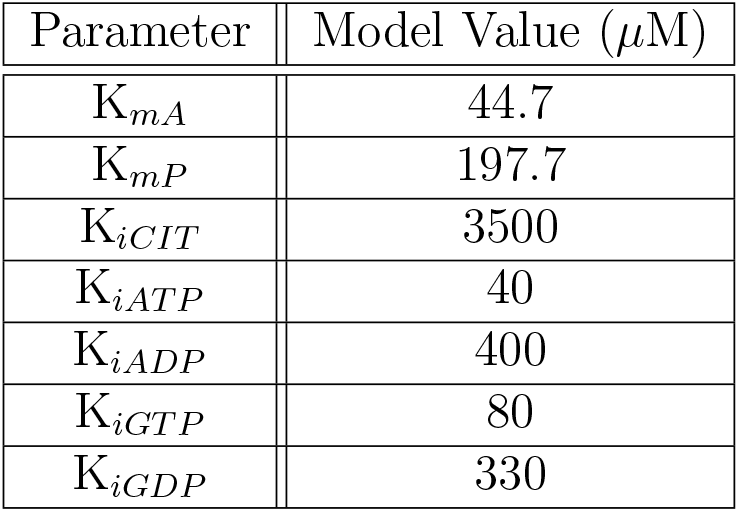
Kinetic parameters for fumarase.

Malate from the previous reaction is consumed by malate dehydrogenase, generating oxaloacetate, which will be used to generate citrate, completing the TCA cycle. Malate dehydrogenase reduces NAD^+^ to NADH during the process. Malate dehydrogenase is competitively inhibited at the NAD^+^/NADH site by adenine nucleotides. Malate dehydrogenase’s kinetics may be described as follows:

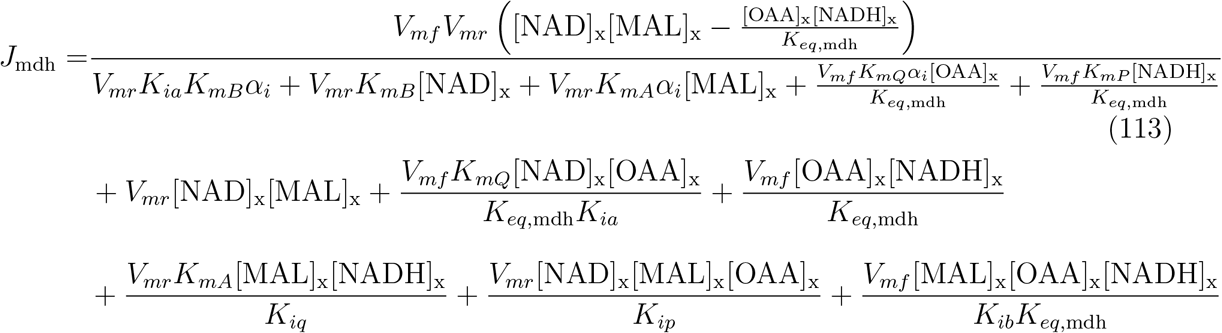

The competitive inhibition by adenine nucleotides is included in the above expression in the *α_i_* term. It is written as follows:

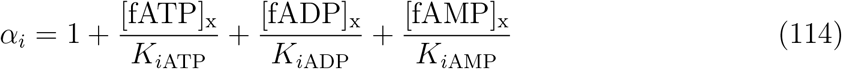

Malate dehydrogenase completes the essential components of the TCA cycle, the reaction mechanism is noted in Wu et al [10]. The kinetic parameters included are listed in Table 9. *V_mr_* may be calculated as 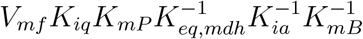.

**Table 9:** Kinetic parameters for malate dehydrogenase.

| Parameter | Model Value ($\mu\text{M}$) |
| --- | --- |
| $K_{mA}$ | 90.55 |
| $K_{mB}$ | 250 |
| $K_{mP}$ | 6.128 |
| $K_{mQ}$ | 2.58 |
| $K_{ia}$ | 279 |
| $K_{ib}$ | 360 |
| $K_{ip}$ | 5.5 |
| $K_{iq}$ | 3.18 |
| $K_{iATP}$ | 183.2 |
| $K_{iADP}$ | 394.4 |
| $K_{iAMP}$ | 420.0 |

We include multiple other reactions, we start with nucleoside diphosphokinase. Nucleoside disphosphokinase transfers a phosphate group from magnesium-bound GTP to a magnesium-bound ADP to produce ATP, we represent its kinetics as follows (see Wu et al. [10] for the reaction mechanism):

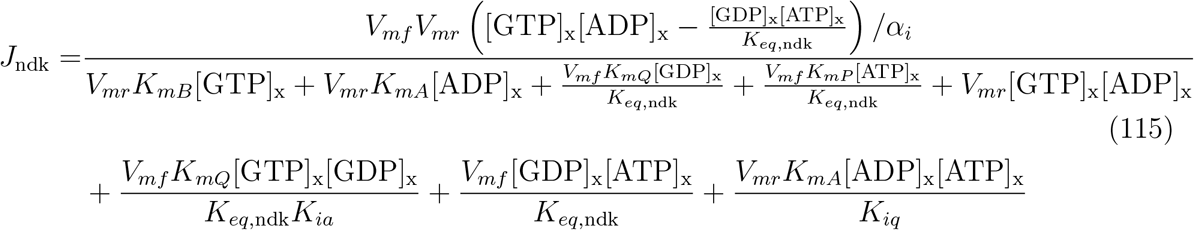

Nucleoside disphosphokinase is regulated by free AMP, leading to the *α_i_* in the above equation, which may be written as follows:

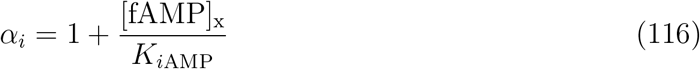

The kinetic parameters included are listed in Table 10. *V_mr_* may be calculated as 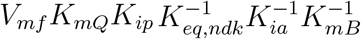.

**Table 10:** Kinetic parameters for nucleoside disphosphokinase.

| Parameter | Model Value ($\mu\text{M}$) |
| --- | --- |
| $K_{mA}$ | 111 |
| $K_{mB}$ | 100 |
| $K_{mP}$ | 260 |
| $K_{mQ}$ | 278 |
| $K_{ia}$ | 170 |
| $K_{ib}$ | 143.6 |
| $K_{ip}$ | 146.6 |
| $K_{iq}$ | 156.5 |
| $K_{iAMP}$ | 650 |

There are only two matrix compounds included in our model that we have not yet discussed, glutamate and aspartate. Glutamate and aspartate are both amino acids, and are both used to a limited extent as a metabolic intermediate. Additionally, glutamate is a neurotransmitter. Glutamate oxaloacetate transaminase (also known as aspartate transaminase) converts aspartate to glutamate, and in the process produces alpha-ketogluterate from oxaloacetate. Glutamate oxaloacetate transaminase’s aspartate binding site is competitively inhibited by alpha-ketogluterate. The reaction mechanism is discussed further in Wu et al [10]:

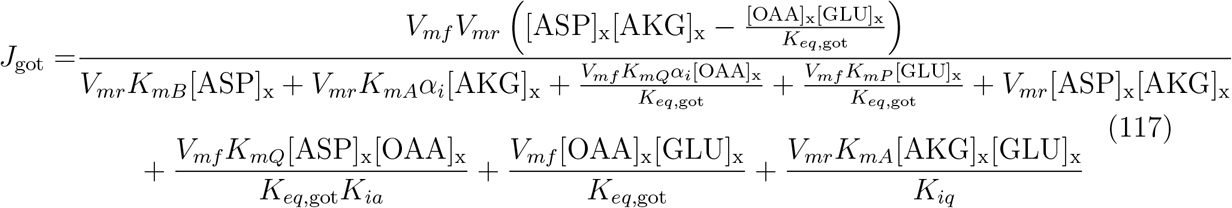

Where *α_i_*, may be expressed as follows:

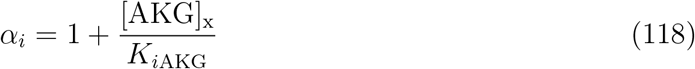

The kinetic parameters included are listed in Table 11. *V_mr_* may be calculated as 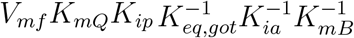.

**Table 11:**
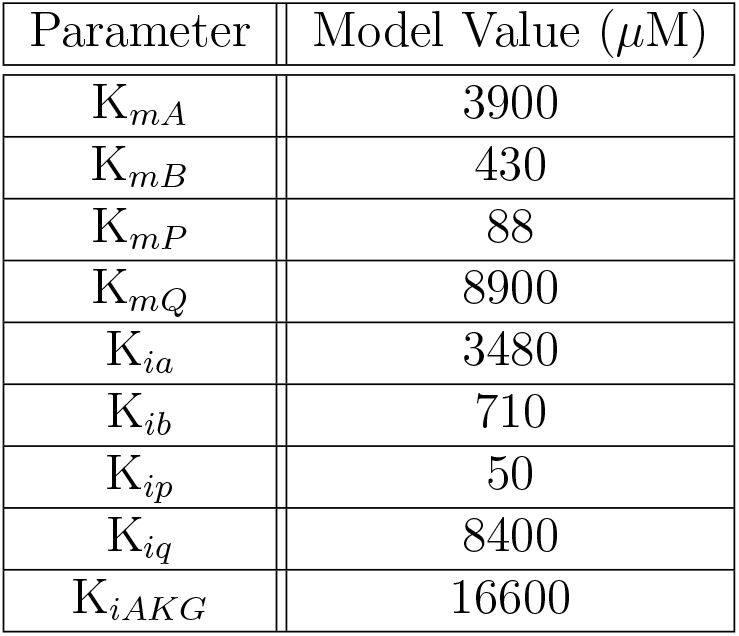
Kinetic parameters for glutamate oxaloacetate transaminase.

| Parameter | Model Value ($\mu\text{M}$) |
| --- | --- |
| $K_{mA}$ | 3900 |
| $K_{mB}$ | 430 |
| $K_{mP}$ | 88 |
| $K_{mQ}$ | 8900 |
| $K_{ia}$ | 3480 |
| $K_{ib}$ | 710 |
| $K_{ip}$ | 50 |
| $K_{iq}$ | 8400 |
| $K_{iAKG}$ | 16600 |

Glutamate may also be consumed directly to produce alpha-ketogluterate by glutamate dehydrogenase. Glutamate dehydrogenase activity is particularly relevant to hepatocytes in the liver [8]. We use a simple kinetic description of glutamate dehydrogenase because the only measurements we could find of glutamate dehydrogenase kinetics, based on the kinetic model used in the experiments we relied on [8]. We represent glutamate dehydrogenase activity as follows:

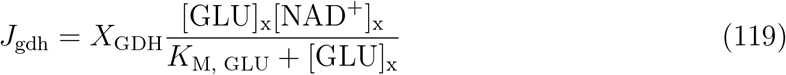

In Jonker et al. [8], they use an even simpler model that did not depend on NAD^+^. We chose *X*_GDH_ so that *V*_GDH_[NAD^+^] was what Jonker et al. measured to be the *V*_max_ of their simpler equation. This change was introduced in order to avoid failures of non-negativity in our model.

Adenylate kinase acts both in the mitochondria and in the cytosol. Adenylate kinase regulates the quantity of AMP in the cell by converting two ADP into AMP and ATP, and is fast enough to be maintained at equilibrium. AMP plays an important regulatory role for cellular metabolism. The kinetics of adenylate kinase are as follows:

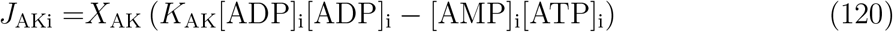

### 1.2 Oxidative Phosphorylation

We will go through the process of oxidative phosphorylation in order, which is particularly crucial to the work that follows. The flux through Complex I depends both on chemical kinetics and the thermodynamic dependence of Complex I activity on the electrical potential gradient:

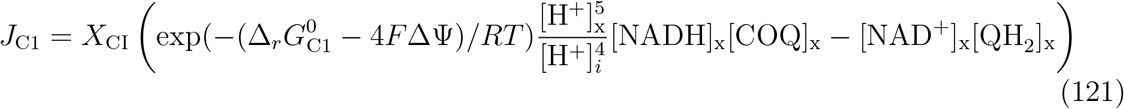

We see that the thermodynamic term gives an exponential dependence of the flux on the electrical potential gradient. The reaction is written so that there is proportional dependence of the forward and backward reactions on their reactants.

Complex III flux has similar kinetics, including exponential dependence on the electrical potential gradient. It primarily differs in the stoichiometry of proton movement via Complex III:

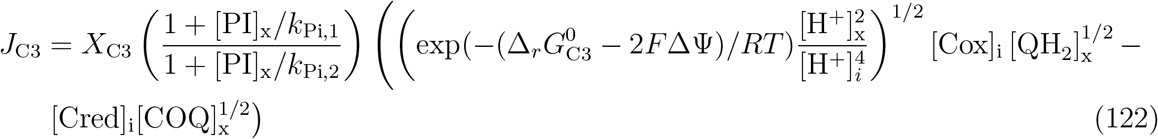

The flux via Complex IV is different because it is not transferring an electron from one intermediate to another, rather it is transferring an electron to its final resting place: an oxygen molecule, in the process producing water. Once again the reaction exhibits exponential dependence on the electrical potential gradient.

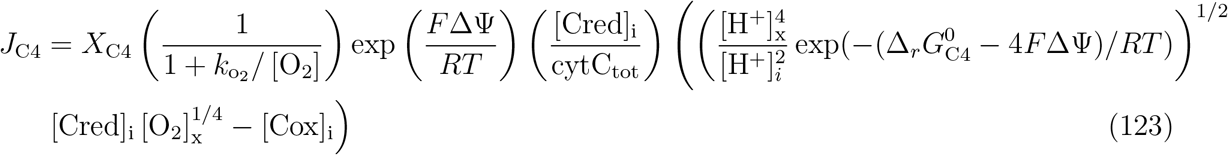

The final step of oxidative phosphorylation is ATP synthase (also known as Complex V or F_0_F_1_-ATPase). ATP Synthase generates a significant portion of the total ATP produced by mitochondrial respiration and consumes the majority of the oxygen used in aerobic respiration:

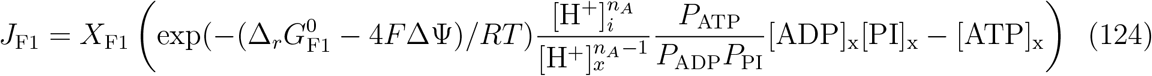

where *P*_ATP_, *P*_ADP_, and *P*_PI_ are the reaction polynomials associated with ATP, ADP, and phosphate respectively. The reaction polynomials may be written explicitly as follows (they’re also discussed above in relation to ion concentrations):

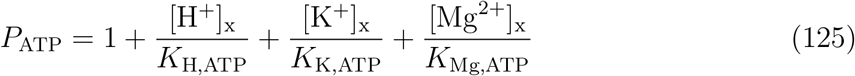

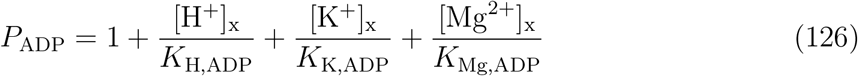

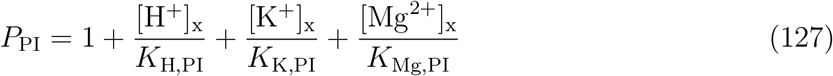

### 1.3 Other Membrane Fluxes

The mitochondrion is composed of two compartments that allow for the reactions involved in cellular respiration to be performed more efficiently as discussed above. The inner membrane is highly impermeable and so relies heavily on transporters to regulate crucial concentrations in the mitochondrion. We will begin our discussion of membrane fluxes with those transporters.

A key transporter for understanding the behaviour of mitochondria is adenine nucleotide translocase. Adenine nucleotide translocase swaps ATP and ADP across the inner mitochondrial membrane. Since ATP synthase converts ADP into ATP in the mitochondrial matrix, the mitochondrial matrix needs a constant supply of ADP and a means of exporting its ATP. More importantly however, adenine nucleotide translocase is a limiting flux for aerobic respiration, meaning that the cell is highly sensitive to the activity of adenine nucleotide translocase. We represent the kinetics of this crucial transporter as follows:

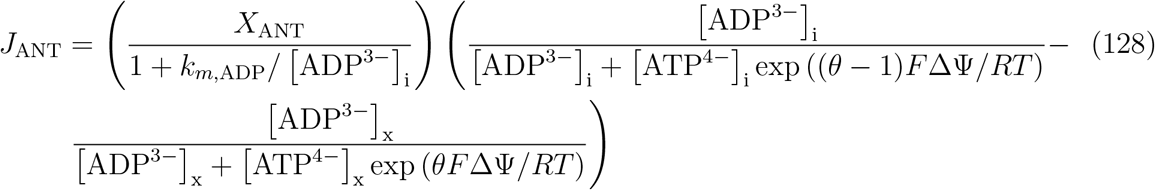

Adenine nucleotide translocase is dependent on the electrical potential gradient because ATP and ADP are both charged, but their exchange is not electroneutral.

Dihydrogen phosphate and a proton are co-transported across the inner membrane of the mitochondrion. This supplies inorganic phosphate ions that may be used in ATP synthesis:

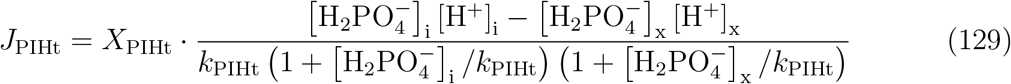

Another crucial transporter is the potassium-hydrogen antiporter. The potassium-hydrogen antiporter exchanges potassium ions for hydrogen ions according to simple mass-action kinetics. Typically hydrogen ions move into the mitochondrial matrix and potassium ions into the intermembrane space. Once again, this flux is electoneutral. This gives us the following kinetics:

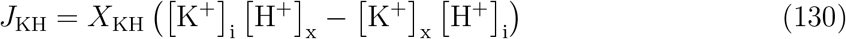

Pyruvate and hydrogen ions are co-transported across the inner membrane of the mitochondrion. Since pyruvate is consumed by pyruvate dehydrogenase inside the cell, the pyruvate-hydrogen co-transporter preferably transports pyruvate and hydrogen into the mitochondrial matrix. Pyruvate-hydrogen co-transporter also follows mass-action kinetics and is electroneutral, giving us the following kinetics:

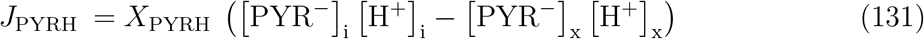

Glutamate and hydrogen ions are also co-transported across the inner membrane of the mitochondrion. When glutamate dehydrogenase activity is high, glutamate will be consumed in the mitochondrial matrix. Otherwise it may not be so clear what flux to expect through the glutamate-hydrogen co-transporter.

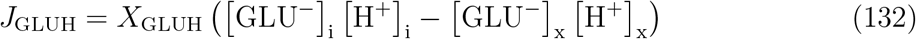

The above two transporters are examples of co-transporters of elemental ions and important metabolites. They provide essential metabolites that the mitochondrion will use. On the other hand, antiporting mechanisms for pairs of metabolites help to maintain appropriate mixes of those pairs of metabolites. Our model includes four examples of this form: citrate-malate antiporter which follows simple mass-action kinetics, alphaketogluterate-malate antiporter, aspartate-glutamate antiporter, and succinate-malate antiporter. The latter three follow more complex kinetics discussed in Wu et al [10]:

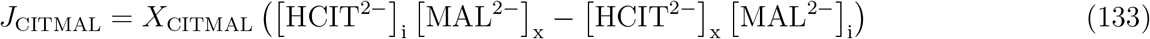

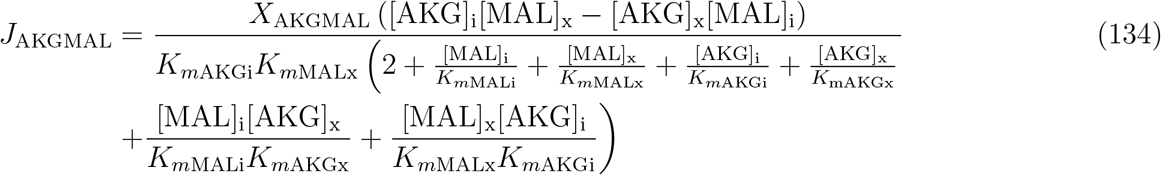

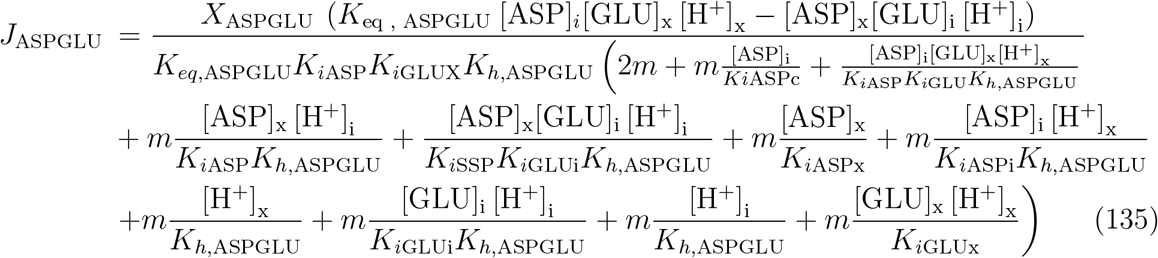

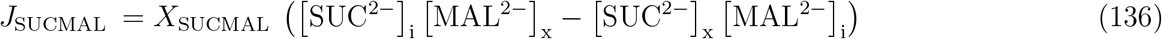

Another regulator of malate concentrations in the mitochondrial matrix is the Malate-Phosphate antiporter. This antiporter follows simple mass-action kinetics, like the citrate-malate antiporter, and it exchanges inorganic phosphate for malate.

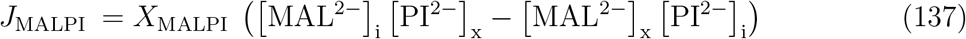

The last pair of fluxes between the mitochondrial matrix and the cell are particularly important, they are the hydrogen and potassium leaks. These effects it is important to mention are not electroneutral, they lead to a net flow of hydrogen and potassium ions into the mitochondrial matrix. The mitochondrial electrical potential gradient will always favour leak into the matrix for the same reason. As noted above in Section ??, these leak fluxes were adjusted for Edwards, Palm, and Layton’s model [5] to match more recent kinetic data on ion leakage across the mitochondria’s inner membrane.

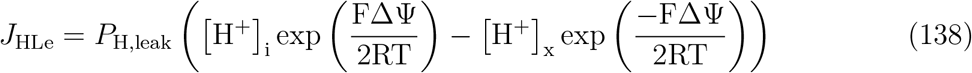

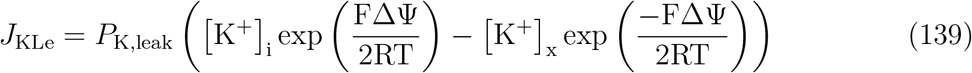

The remaining membrane fluxes of interest connect the cytosol and the intermembrane space. These fluxes represent passive diffusion because the outer mitochondrial membrane is highly permeable and doesn’t for the most part depend on transporters to control the flux in the quantities we include. The passive diffusion fluxes we include are as follows:

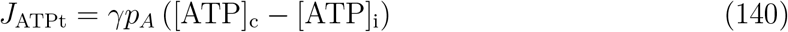

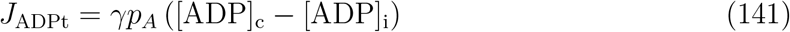

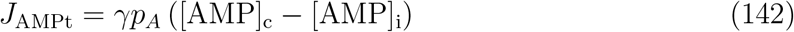

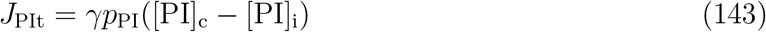

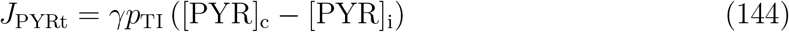

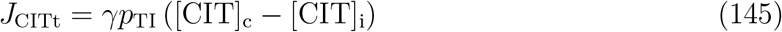

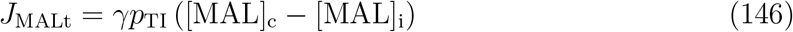

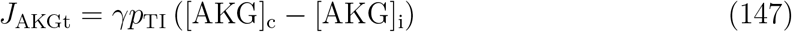

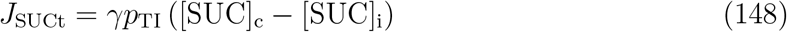

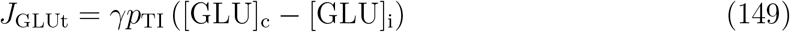

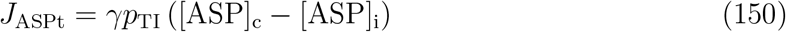

The outer membrane is highly permeable and so the cytosol and intermembrane space typically have very similar concentrations of all the above compounds.

### 1.4 Other Reactions in the Cytosol

ATP consumption is the most important remaining flux in the model. ATP consumption is treated as independent of the system’s state by the previous model [5]. However we use Michaelis-Menten dynamics for the cell’s ATP consumption (*Q*_ATP_), i.e.

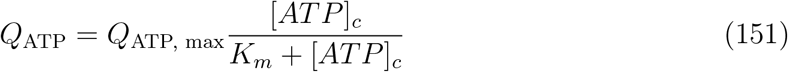

The parameterization of this equation is discussed in Section ??.

We also include glycolysis in some tissues (the liver by default, the mTAL optionally) as follows (with *K_mGlyc_* chosen to be so small that it only serves to ensure non-negativity of ADP levels for an otherwise relatively constant source term):

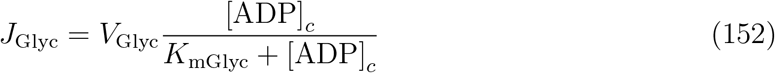

In the cytosol, adenylate kinase (represented in our model by *J*_AKc_) and creatine kinase (represented in our model by *J*_CKc_) impact adenine nucleotide homeostasis by controlling AMP concentrations. Aside from being an adenine nucleotide, AMP plays an important regulatory role in the cell. These reactions have a high capacity in our model and are kept near equilibrium:

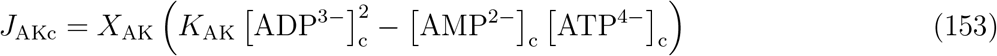

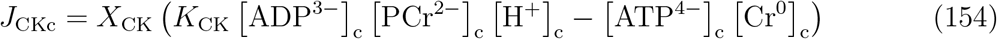

These reactions exhaust the fluxes included in our model.

### 1.5 Full List of State Variables

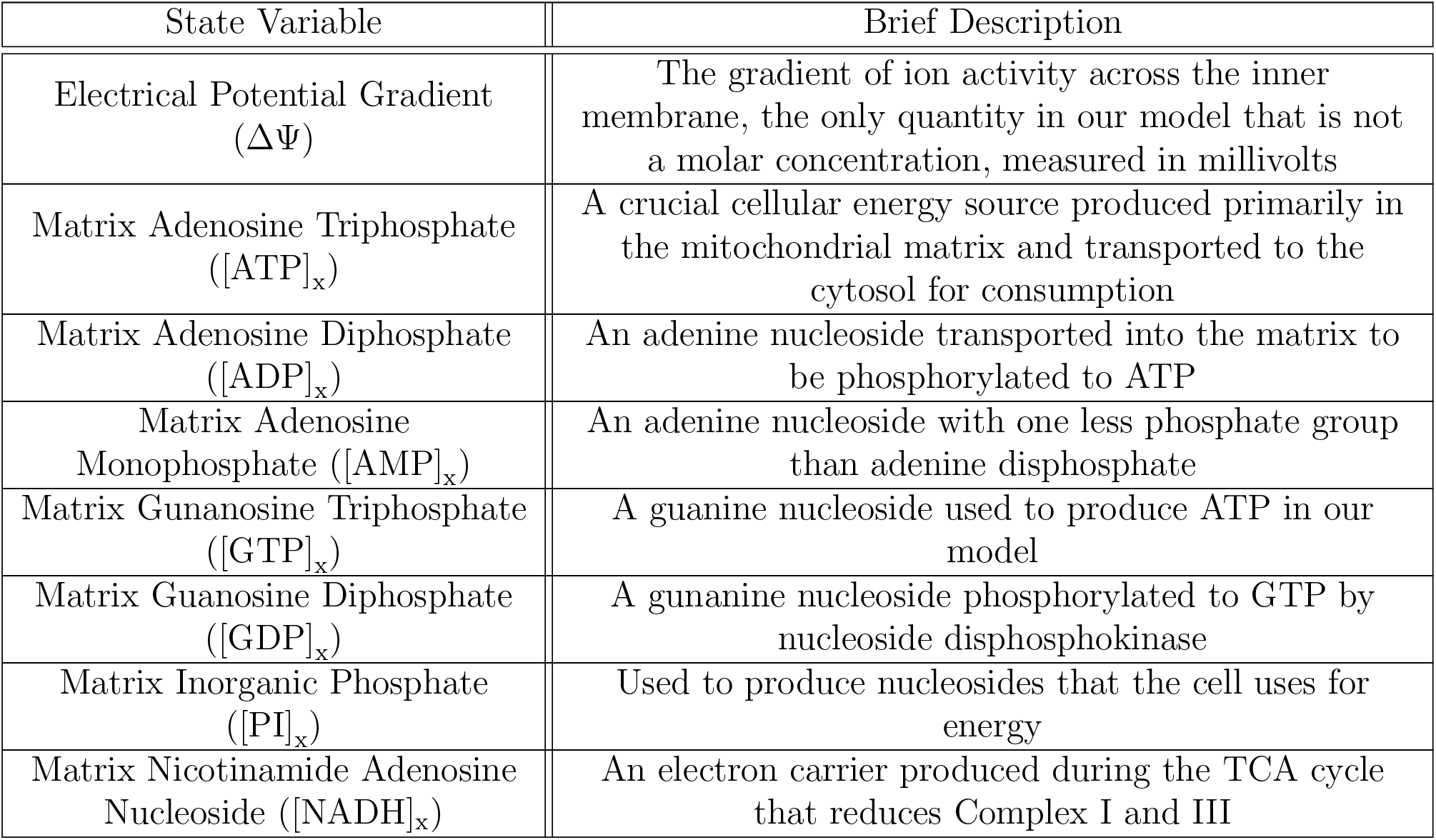
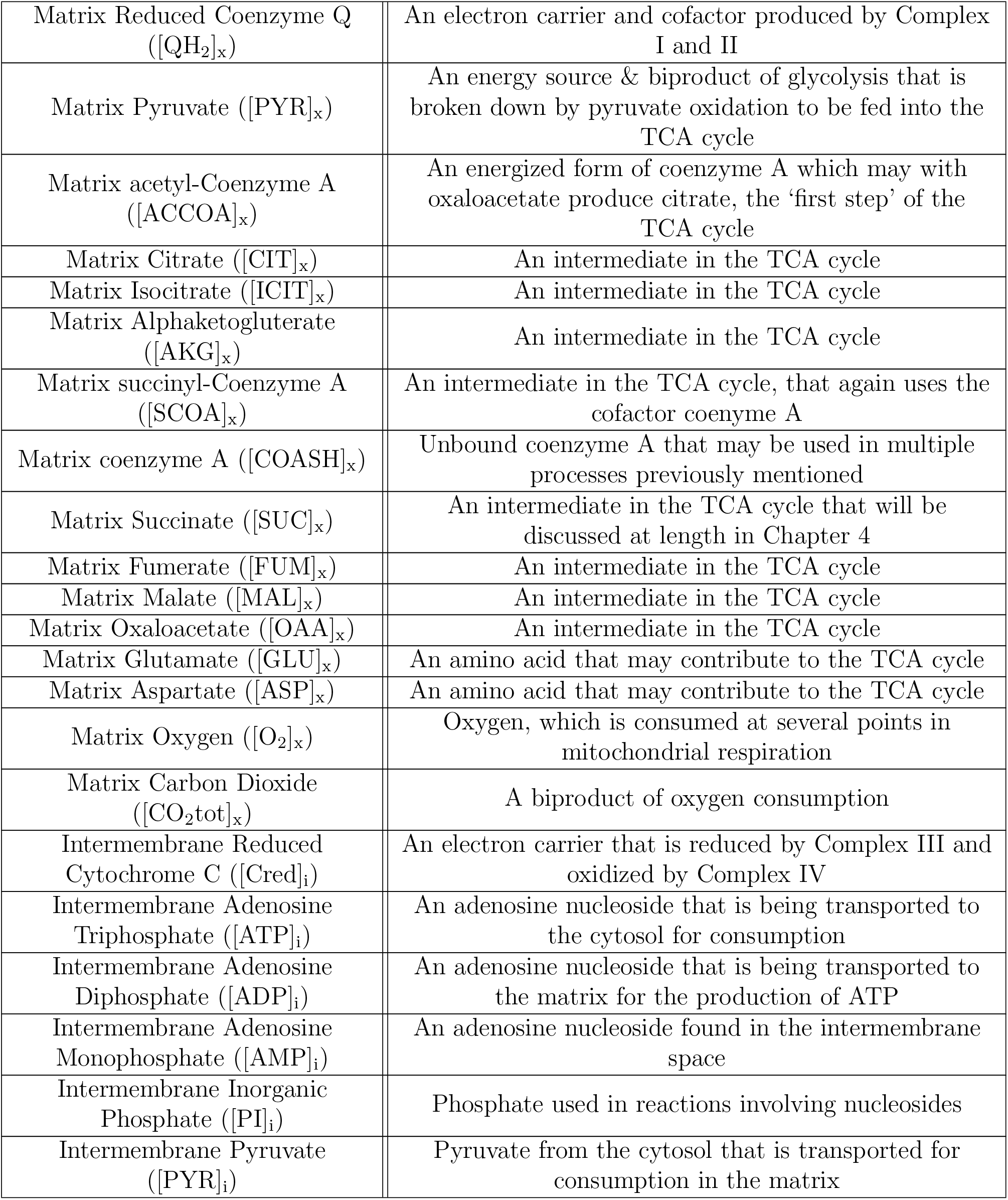
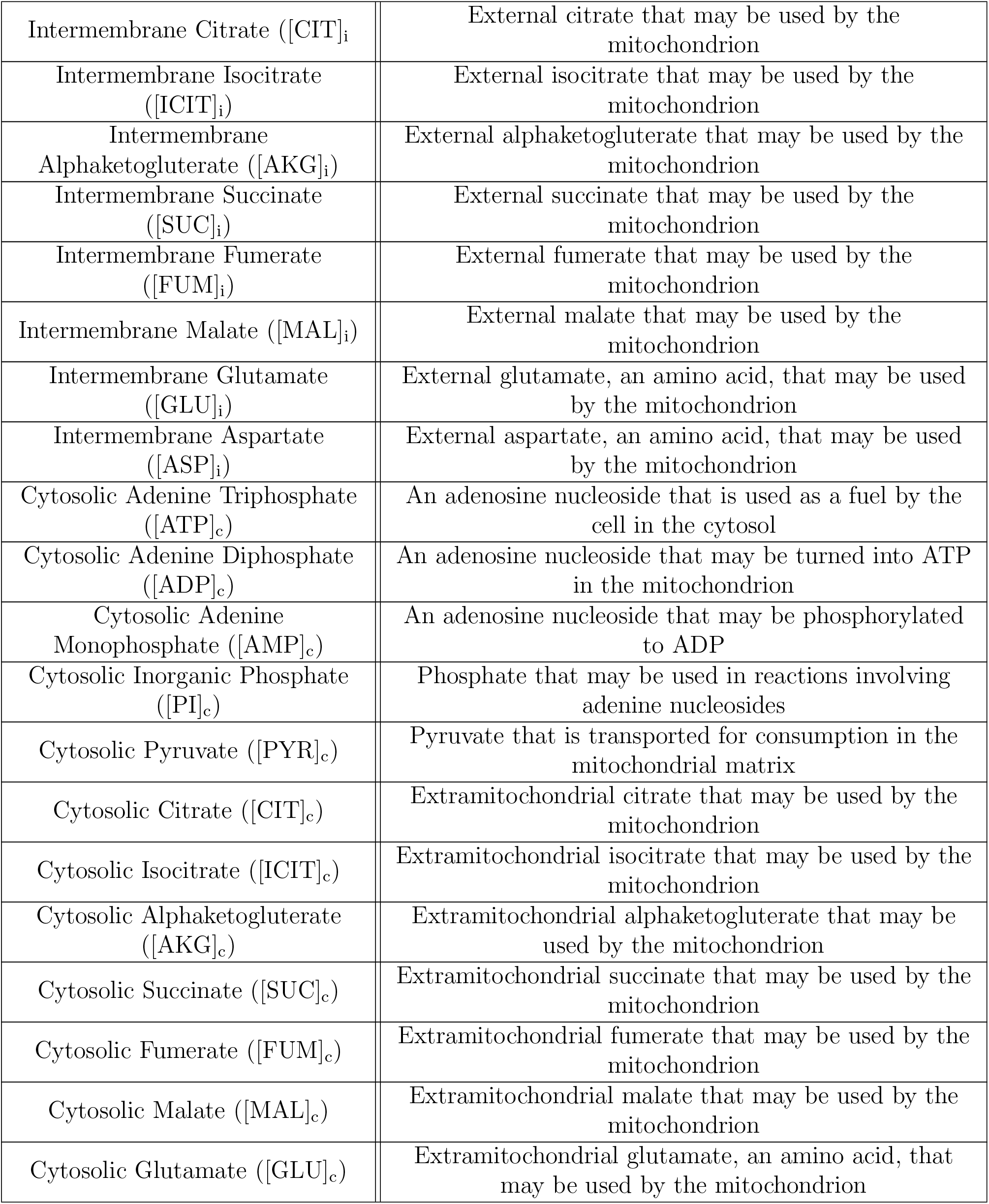
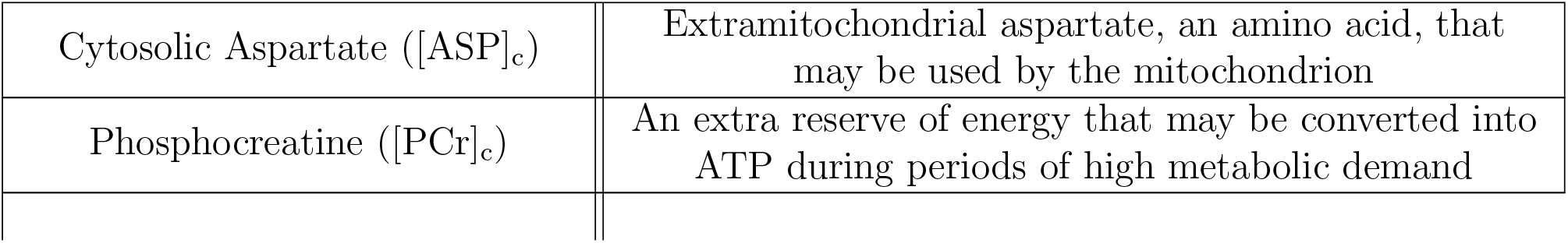

| State Variable | Brief Description |
| --- | --- |
| Electrical Potential Gradient<br>($\Delta\Psi$) | The gradient of ion activity across the inner membrane, the only quantity in our model that is not a molar concentration, measured in millivolts |
| Matrix Adenosine Triphosphate<br>($[ATP]_x$) | A crucial cellular energy source produced primarily in the mitochondrial matrix and transported to the cytosol for consumption |
| Matrix Adenosine Diphosphate<br>($[ADP]_x$) | An adenine nucleoside transported into the matrix to be phosphorylated to ATP |
| Matrix Adenosine Monophosphate<br>($[AMP]_x$) | An adenine nucleoside with one less phosphate group than adenine diphosphate |
| Matrix Guanosine Triphosphate<br>($[GTP]_x$) | A guanine nucleoside used to produce ATP in our model |
| Matrix Guanosine Diphosphate<br>($[GDP]_x$) | A guanine nucleoside phosphorylated to GTP by nucleoside diphosphokinase |
| Matrix Inorganic Phosphate<br>($[PI]_x$) | Used to produce nucleosides that the cell uses for energy |
| Matrix Nicotinamide Adenosine Nucleoside<br>($[NADH]_x$) | An electron carrier produced during the TCA cycle that reduces Complex I and III |
| Matrix Reduced Coenzyme Q ([QH<sub>2</sub>]<sub>x</sub>) | An electron carrier and cofactor produced by Complex I and II |
| Matrix Pyruvate ([PYR]<sub>x</sub>) | An energy source & biproduct of glycolysis that is broken down by pyruvate oxidation to be fed into the TCA cycle |
| Matrix acetyl-Coenzyme A ([ACCOA]<sub>x</sub>) | An energized form of coenzyme A which may with oxaloacetate produce citrate, the ‘first step’ of the TCA cycle |
| Matrix Citrate ([CIT]<sub>x</sub>) | An intermediate in the TCA cycle |
| Matrix Isocitrate ([ICIT]<sub>x</sub>) | An intermediate in the TCA cycle |
| Matrix Alphaketogluterate ([AKG]<sub>x</sub>) | An intermediate in the TCA cycle |
| Matrix succinyl-Coenzyme A ([SCOA]<sub>x</sub>) | An intermediate in the TCA cycle, that again uses the cofactor coenzyme A |
| Matrix coenzyme A ([COASH]<sub>x</sub>) | Unbound coenzyme A that may be used in multiple processes previously mentioned |
| Matrix Succinate ([SUC]<sub>x</sub>) | An intermediate in the TCA cycle that will be discussed at length in Chapter 4 |
| Matrix Fumerate ([FUM]<sub>x</sub>) | An intermediate in the TCA cycle |
| Matrix Malate ([MAL]<sub>x</sub>) | An intermediate in the TCA cycle |
| Matrix Oxaloacetate ([OAA]<sub>x</sub>) | An intermediate in the TCA cycle |
| Matrix Glutamate ([GLU]<sub>x</sub>) | An amino acid that may contribute to the TCA cycle |
| Matrix Aspartate ([ASP]<sub>x</sub>) | An amino acid that may contribute to the TCA cycle |
| Matrix Oxygen ([O<sub>2</sub>]<sub>x</sub>) | Oxygen, which is consumed at several points in mitochondrial respiration |
| Matrix Carbon Dioxide ([CO<sub>2</sub>tot]<sub>x</sub>) | A biproduct of oxygen consumption |
| Intermembrane Reduced Cytochrome C ([Cred]<sub>i</sub>) | An electron carrier that is reduced by Complex III and oxidized by Complex IV |
| Intermembrane Adenosine Triphosphate ([ATP]<sub>i</sub>) | An adenosine nucleoside that is being transported to the cytosol for consumption |
| Intermembrane Adenosine Diphosphate ([ADP]<sub>i</sub>) | An adenosine nucleoside that is being transported to the matrix for the production of ATP |
| Intermembrane Adenosine Monophosphate ([AMP]<sub>i</sub>) | An adenosine nucleoside found in the intermembrane space |
| Intermembrane Inorganic Phosphate ([PI]<sub>i</sub>) | Phosphate used in reactions involving nucleosides |
| Intermembrane Pyruvate ([PYR]<sub>i</sub>) | Pyruvate from the cytosol that is transported for consumption in the matrix |
| Intermembrane Citrate ([CIT]<sub>i</sub>) | External citrate that may be used by the mitochondrion |
| Intermembrane Isocitrate ([ICIT]<sub>i</sub>) | External isocitrate that may be used by the mitochondrion |
| Intermembrane Alphaketoglutarate ([AKG]<sub>i</sub>) | External alphaketoglutarate that may be used by the mitochondrion |
| Intermembrane Succinate ([SUC]<sub>i</sub>) | External succinate that may be used by the mitochondrion |
| Intermembrane Fumerate ([FUM]<sub>i</sub>) | External fumerate that may be used by the mitochondrion |
| Intermembrane Malate ([MAL]<sub>i</sub>) | External malate that may be used by the mitochondrion |
| Intermembrane Glutamate ([GLU]<sub>i</sub>) | External glutamate, an amino acid, that may be used by the mitochondrion |
| Intermembrane Aspartate ([ASP]<sub>i</sub>) | External aspartate, an amino acid, that may be used by the mitochondrion |
| Cytosolic Adenine Triphosphate ([ATP]<sub>c</sub>) | An adenosine nucleoside that is used as a fuel by the cell in the cytosol |
| Cytosolic Adenine Diphosphate ([ADP]<sub>c</sub>) | An adenosine nucleoside that may be turned into ATP in the mitochondrion |
| Cytosolic Adenine Monophosphate ([AMP]<sub>c</sub>) | An adenosine nucleoside that may be phosphorylated to ADP |
| Cytosolic Inorganic Phosphate ([PI]<sub>c</sub>) | Phosphate that may be used in reactions involving adenine nucleosides |
| Cytosolic Pyruvate ([PYR]<sub>c</sub>) | Pyruvate that is transported for consumption in the mitochondrial matrix |
| Cytosolic Citrate ([CIT]<sub>c</sub>) | Extramitochondrial citrate that may be used by the mitochondrion |
| Cytosolic Isocitrate ([ICIT]<sub>c</sub>) | Extramitochondrial isocitrate that may be used by the mitochondrion |
| Cytosolic Alphaketoglutarate ([AKG]<sub>c</sub>) | Extramitochondrial alphaketoglutarate that may be used by the mitochondrion |
| Cytosolic Succinate ([SUC]<sub>c</sub>) | Extramitochondrial succinate that may be used by the mitochondrion |
| Cytosolic Fumerate ([FUM]<sub>c</sub>) | Extramitochondrial fumerate that may be used by the mitochondrion |
| Cytosolic Malate ([MAL]<sub>c</sub>) | Extramitochondrial malate that may be used by the mitochondrion |
| Cytosolic Glutamate ([GLU]<sub>c</sub>) | Extramitochondrial glutamate, an amino acid, that may be used by the mitochondrion |
| Cytosolic Aspartate ([ASP]<sub>c</sub>) | Extramitochondrial aspartate, an amino acid, that may be used by the mitochondrion |
| Phosphocreatine ([PCr]<sub>c</sub>) | An extra reserve of energy that may be converted into ATP during periods of high metabolic demand |

### 1.6 List of Significant Parameters

**Table 13:** Adjustable parameters of the model that are fitted in previous work or estimated below [10]. In each reaction expression where there is a *V_mf_* and *V_mr_* term, the *V_mf_* (or forward reaction) term is the associated activity, and *V_mr_* may be calculated as explained in the relevant section above.

| Parameter | Units | Brief Description |
| --- | --- | --- |
| $X_{PDH}$ | $M^4/s$ | Pyruvate dehydrogenase activity |
| $X_{CITS}$ | $M/s$ | Citrate synthase activity |
| $X_{ACON}$ | $M/s$ | Acontinase activity |
| $X_{ISOD}$ | $M/s$ | Isocitrate dehydrogenase activity |
| $X_{AKGD}$ | $M/s$ | Alphaketoglutarate dehydrogenase activity |
| $X_{SCOAS}$ | $M/s$ | Succinyl-CoA synthetase activity |
| $X_{SDH}$ | $M/s$ | Succinate dehydrogenase activity |
| $X_{FUM}$ | $M/s$ | Fumarase activity |
| $X_{MDH}$ | $M/s$ | Malate dehydrogenase activity |
| $X_{NDK}$ | $M/s$ | Nucleoside diphosphokinase activity |
| $X_{GOT}$ | $M/s$ | Glutamaete oxaloacetate transaminase activity |
| $X_{GDH}$ | $s^{-1}$ | Glutamate dehydrogenase activity |
| $X_{PYRH}$ | $M/s$ | Pyruvate-hydrogen cotransporter activity |
| $X_{GLUH}$ | $M/s$ | Glutamate-hydrogen cotransporter activity |
| $X_{CITMAL}$ | $M/s$ | Citrate-Malate exchange transporter activity |
| $X_{AKGMAL}$ | $M/s$ | Alphaketoglutarate-Malate exchange transporter activity |
| $X_{MALPI}$ | $M/s$ | Malate-Phosphate exchange transporter activity |
| $X_{ASPGLU}$ | $M/s$ | Aspartate-Glutamate exchange transporter activity |
| $X_{SUCMAL}$ | $M/s$ | Succinate-Malate exchange transporter activity |
| $X_{HK}$ | $M/s$ | Hexokinase activity |
| $X_{CI}$ | $M/s$ | Complex I activity |
| $X_{CIII}$ | $M/s$ | Complex III activity |
| $X_{CIV}$ | $M/s$ | Complex IV activity |
| $X_{ATP}$ | $M/s$ | Complex V/ATPase/ATP Synthase activity |
| $X_{ANT}$ | $M/s$ | Adenine nucleotide translocase activity |
| $X_{PIHt}$ | $M/s$ | Phosphate-hydrogen co-transporter activity |
| $k_{PiH}$ | $M$ | Phosphate-hydrogen regulative parameter |
| $X_{KH}$ | $M/s$ | Potassium-hydrogen antiport activity |
| $P_{H, leak}$ | $s^{-1}$ | Hydrogen leak permeability |
| $k_{Pi,1}$ | $M$ | Determines the dependence of Complex III on phosphate, producing a more sensitive dependence |
| $k_{Pi,2}$ | $M$ | Determines the dependence of Complex III on phosphate, producing a less sensitive dependence |
| $W_x$ | (L water)/(L matrix) | Mitochondrial matrix water fraction |
| $W_i$ | (L water)/(L intermembrane space) | Mitochondrial intermembrane space water fraction |
| $W_c$ | (L water)/(L cytoplasm) | Cytosolic water fraction |
| $C_{tot}$ | M | Total concentration of reduced and oxidized cytochrome C in intermembrane space |
| $Q_{tot}$ | M | Total concentration of reduced and oxidized coenzyme Q in mitochondrial matrix |
| $NAD_{tot}$ | M | Total concentration of reduced and oxidized $NAD^+/NADH$ in mitochondrial matrix |
| $FAD_{tot}$ | M | Total concentration of reduced and oxidized $FAD^{2+}/FADH_2$ |
| $k_{O_2}$ | M | Oxygen affinity of Complex V |
| $k_{mADP}$ | M | ADP affinity of Complex V |
| $P_{K, leak}$ | $s^{-1}$ | Potassium leak permeability |
| $Q_{ATP, max}$ | M/s | Maximal ATP consumption |
| $K_{mATP}$ | M | $K_m$ for ATP consumption |
| $V_{Glyc}$ | M/s | Maximal Glycolytic ATP production |
| $K_{mGlyc}$ | M | $K_m$ for glycolytic ATP production |
| $K_{CK}$ | $M^{-1}$ | Creatine kinase equilibrium constant |
| $\gamma$ | $\mu m^2$ | Outer membrane area per mitochondrial volume |
| $\rho_m$ | (mg prot) / (L mito) | Protein density of mitochondria |
| $V_{mito}$ | (L mito)/(L cell) | Mitochondrial volume fraction |
| $V_{cyto}$ | (L cyto)/(L cell) | Cytoplasmic volume fraction |
| $n_A$ | Unitless | Stoichiometry for hydrogen ion transport by Complex V |
| $p_{PI}$ | $\mu m/s$ | Permeability of outer membrane to inorganic phosphate |
| $p_A$ | $\mu m/s$ | Permeability of outer membrane to nucleotides |
| $k_{m, ADP}$ | M | ANT Michaelis-Menten constant |
| $\theta$ | Unitless | ANT kinetic parameter |
| $\beta$ | M | Matrix buffering capacity |
| $C_{IM}$ | mol/(L mito mV) | Capacitance of inner membrane for electrical potential gradient |
| $B_x$ | M | Matrix buffering parameter |
| $K_{Bx}$ | M | Matrix buffering parameter |

### 1.7 List of Physical Parameters

**Table 14:** Non-adjustable physical constants used in our model.

| Physical Parameter | Value | Units | Brief Description |
| --- | --- | --- | --- |
| RT | 2.5775 | kJ/mol | Gas constant times temperature |
| F | 0.096484 | kJ/(mol mV) | Faraday’s constant |

## 2 Sensitivity Analysis

### 2.1 Local sensitivity analysis

We consider the change in state variables in the proximal tubule under a change in each parameter of our model. The full local sensitivity analysis is reported in Figure 1. Here we focus on several key state variables and the parameters that are most impactful upon these state variables. In Figure 3 we show the derivative of each state variable against each of the most significant parameters, calculated using a central difference scheme with Δ*p* = 0.01*p* where *p* is the size of the parameter. The derivative is normalized relative to the baseline size of each state variable and the size of each parameter. We see that the concentrations of NADH and reduced coenzyme Q (QH_2_) are most sensitive to the parameter changes we considered. The cytosolic ATP concentration is most sensitive to the maximal ATP consumption. The concentration of NADH in the cell is determined heavily by alphaketogluterate dehydrogenase, which is positively associated with increased NADH content, and pyruvate dehydrogenase, which is negatively associated with NADH content. Greater alphaketogluterate dehydrogenase activity allows for more production of NADH because in the proximal tubule it is predicted to be a limiting step for the TCA cycle. Alphaketogluterate concentrations are higher in our model than any other TCA cycle intermediate, and the concentration of the product of alphaketogluterate dehydrogenase, succinyl-CoA, is lower than any TCA cycle intermediate. Pyruvate dehydrogenase on the other hand reduces NADH concentrations because it competes with alphaketogluterate dehydrogenase for coenzyme A as a substrate. Pyruvate dehydrogenase uses coenzyme A to produce acetyl-CoA whereas alphaketogluterate uses coenzyme A to produce succinyl-CoA. In the full sensitivity plot, Figure 1, we see that the TCA cycle intermediates from citrate to alphaketogluterate are present in greater concentrations when you increase pyruvate dehydrogenase activity, and intermediates down-stream of alphaketogluterate dehydrogenase are present in lower quantities. Supplementary simulations found that for 1% or 10% increases in the activity of pyruvate dehydrogenase, we observe a consistent alphaketogluterate dehydrogenase flux, despite larger concentrations of alphaketogluterate. This observation further supports this proposed mechanism, since more alphaketogluterate is necessary to produce the same enzyme flux. Aside from the above mechanism and its downstream effects, we see unsurprisingly that the concentration of reduced coenzyme Q and reduced cytochrome C continues to increase with increased total quantities of coenzyme Q and cytochrome C. We also see that the passing of electrons from coenzyme Q to cytochrome C appears to be limited by the available cytochrome C, since the reduction state of coenzyme Q appears to be highly sensitive to the total cytochrome C concentration.

**Figure 1:**
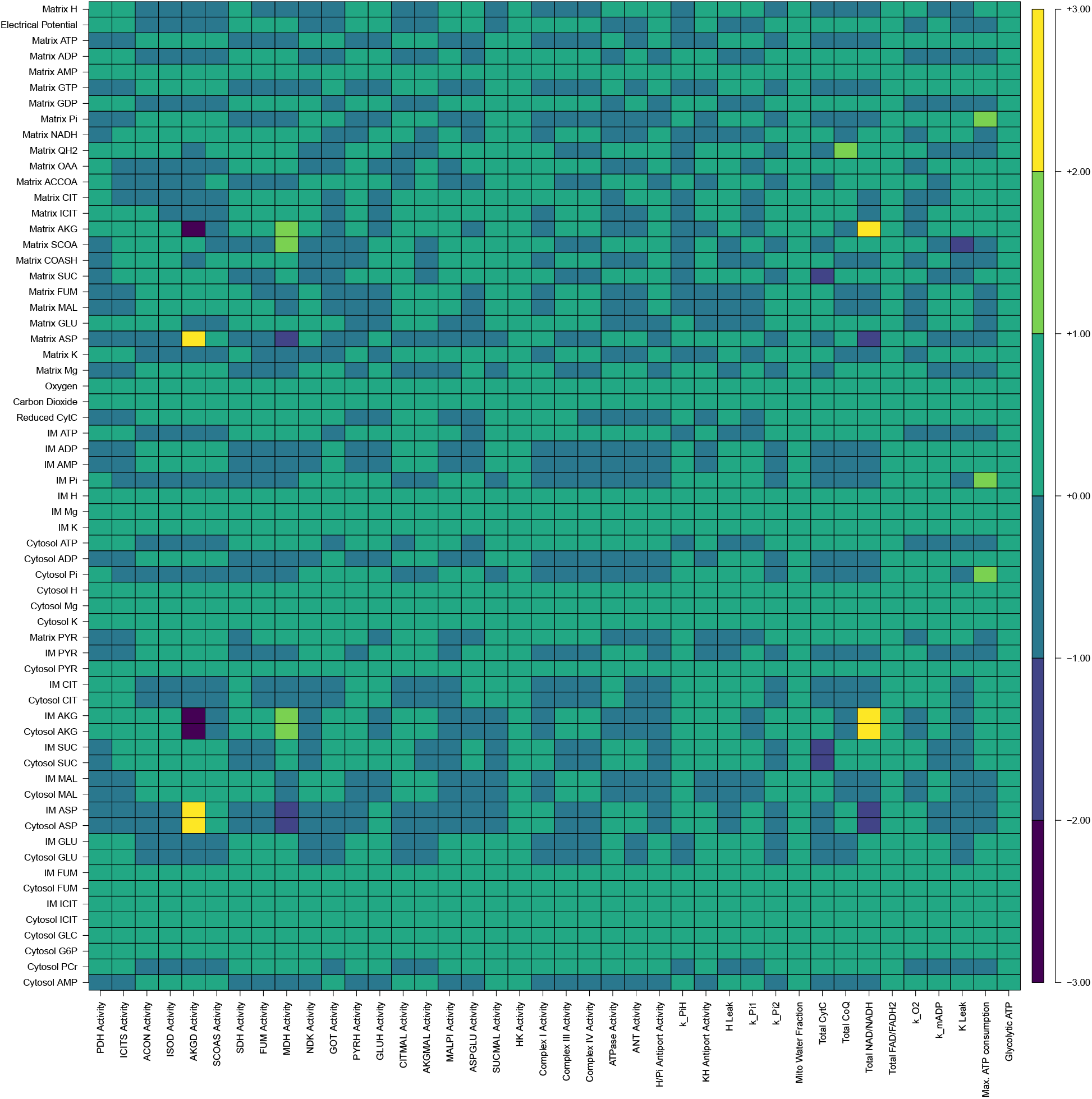
The full set of local sensitivities in the proximal tubule, for more information see section 2.1.

In the medullary thick ascending limb (mTAL) we find several differences and an overall greater robustness to small perturbations of key parameters. The full local sensitivity analysis is reported in Figure 2. In Figure 3 we show the sensitivity calculated in the same manner as for the proximal tubule. At the same threshold for a parameter to be significant, two more parameters are significantly sensitive in the proximal tubule than are sensitive in the mTAL. The concentrations of NADH, reduced coenzyme Q (QH_2_), and reduced cytochrome C are particularly sensitive state variables for the mTAL. The most significant difference between the proximal tubule and mTAL is that in the mTAL, NADH concentrations are less sensitive to pyruvate dehydrogenase and alphaketogluterate dehydrogenase activities. Instead we see in the mTAL that the pooled concentration of NADH and NAD^+^ is by far the most important factor impacting the concentration of NADH. This suggests that whereas in the proximal tubule there are limited amounts of coenzyme A and this limits the amount available to alphaketogluterate dehydrogenase (as illustrated by the sensitivity to pyruvate dehydrogenase activity), instead NAD^+^ is in short supply in the mTAL, and so the way to provide more NADH to the cell is to increase the size of the NADH/NAD^+^ pool. Given the higher NADH to NAD^+^ ratio in the mTAL and its relative hypoxia (which can leave the electron transport chain in a more reduced state [7]), coenzyme Q is also in high demand, leading to a high sensitivity of multiple states to the pooled coenzyme Q concentration.

**Figure 2:**
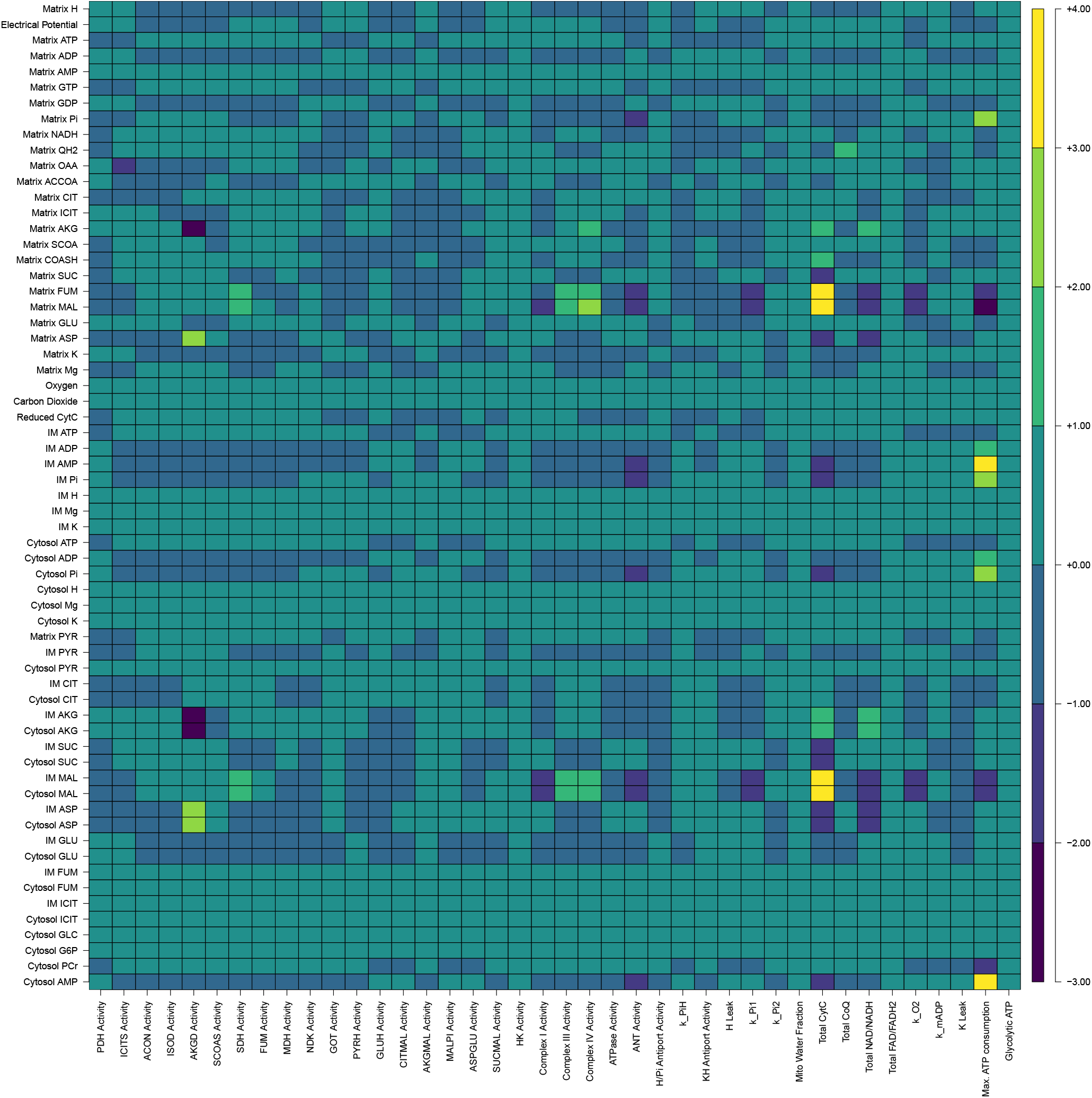
The full set of local sensitivities in the mTAL, for more information see section 2.1.

**Figure 3:**
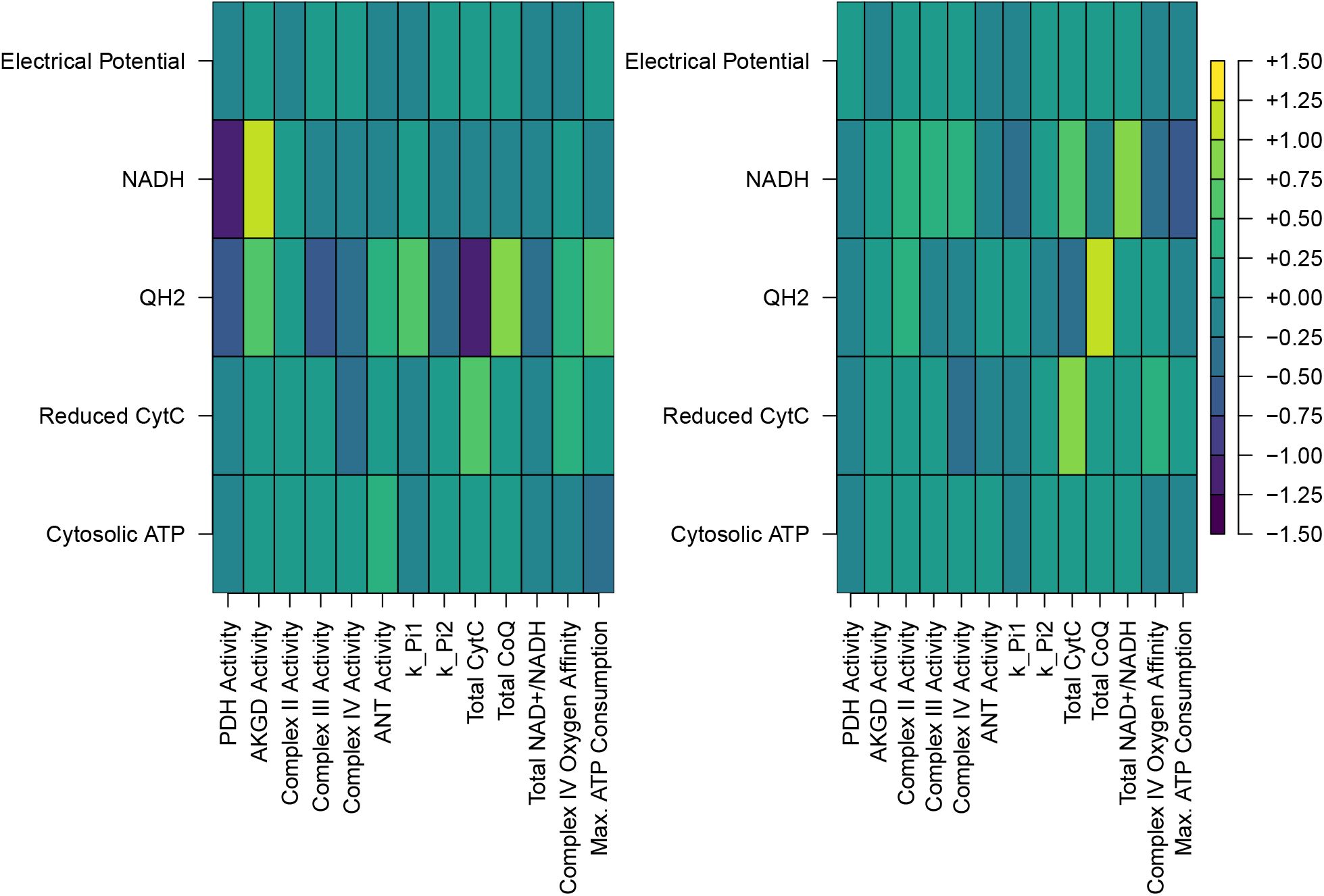
Local sensitivities of several important state variables in the proximal tubule (left panel) and mTAL (right panel) relative to certain parameters significant to either the proximal tubule or mTAL. The results are calculated as described in Section 2.1. CytC stands for cyto chrome C, reduced cytochrome C refers to cytochrome C that has been donated an electron by Complex III. AKGD refers to alphaketogluterate dehydrogenase, PDH refers to pyruvate dehydrogenase, ANT refers to adenine nucleotide translocase, and QH2 is the reduced form of coenzyme Q.

**Figure 4:**
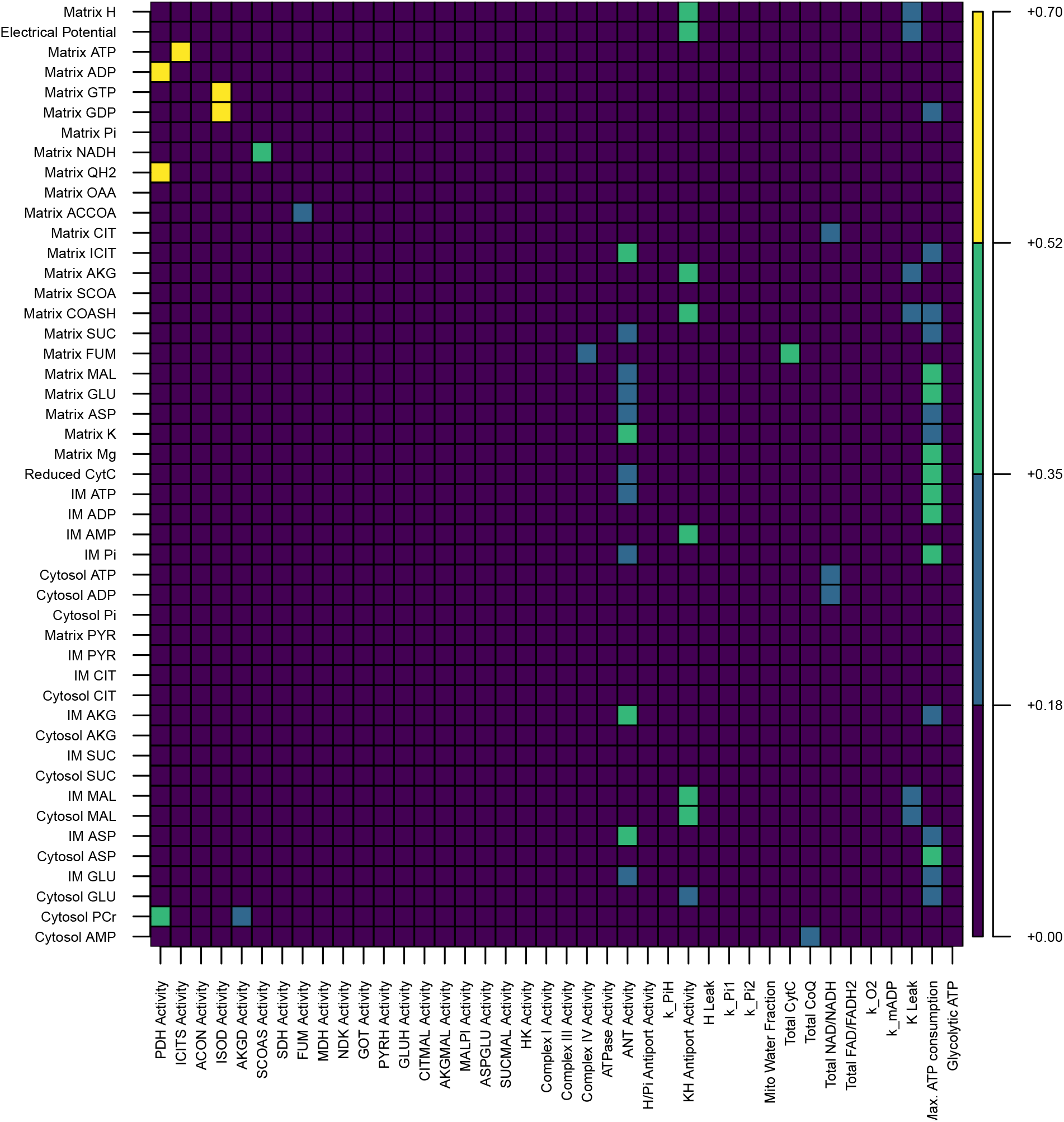
The sensitivity of all state variables to the parameters as calculated according to the Sobol method, without interactions.

**Figure 5:**
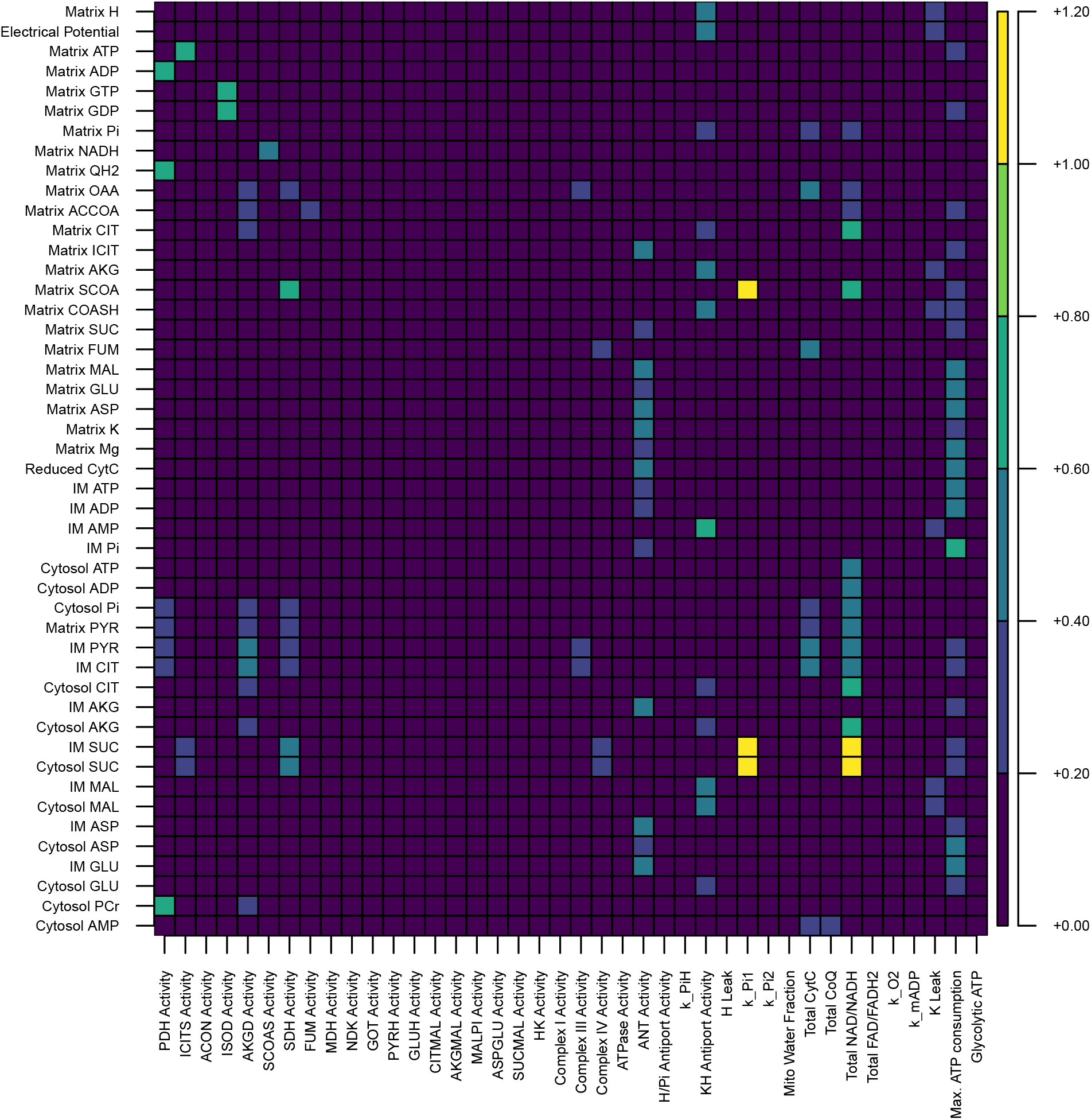
The sensitivity of all state variables to the parameters as calculated according to the Sobol method, taking into account two-way interactions.

### 2.2 Global sensitivity analysis

For our global sensitivity analysis we used the Sobol method for uniformly exploring a highdimensional parameter space and describing the observed variation in the state space as a function of the parameters. For local estimates, interactions are negligible, but in a global sensitivity analysis interactions may be very important. For this reason we report two sensitivity estimates, the first Sobol indices, which do not account for interactions and the total Sobol indices, which account for two-way interactions. First Sobol indices are non-negative numbers between zero and one, whereas total Sobol indices may be greater than one. They represent the proportion of variance explained by a particular parameter in the case of the first Sobol index, and in the case of the total Sobol index, the variance explained by the parameter and the pair-wise interactions associated with it. In Figure 6 we present the first Sobol indices for several important state variables, and in Figure 7 we present the total Sobol indices for those state variables. We see that the first Sobol index results share some features with the results above. We see that like for the local sensitivity analysis of the proximal tubule, pyruvate dehydrogenase features prominently, its importance is discussed above in our local sensitivity analysis. Succinyl-CoA synthetase *in vivo* catalyzes the conversion of succinyl-CoA into succinate and coenzyme A. This may increase the concentration of NADH by increasing the availability of coenzyme A, which in the proximal tubule and mTAL is found in lower concentrations than acetyl-CoA and succinyl-CoA. This could explain its effects on NADH concentrations in the mitochondrion. The next-most important parameter for influencing the state variables is the maximal ATP consumption, which plays a large role in determining the reduced cytochrome C concentration. ATP consumption frees ADP for phosphorylation. This relieves the proton gradient across the inner membrane. Complex IV requires protons in the mitochondrial matrix in order to oxidize cytochrome C. When we consider pairwise interactions via the total Sobol index we see differences in which parameters stand out. Pyruvate dehydrogenase and succinyl-CoA synthetase still impact coenzyme Q and NADH concentrations. Most notably the activities of adenine nucleotide translocase, alphaketogluterate dehydrogenase, and potassium leak are all significant when we consider interactions. Aside from this the results are fairly similar, and we don’t see large changes in the influence of the most impactful parameters on the important state variables.

**Figure 6:**
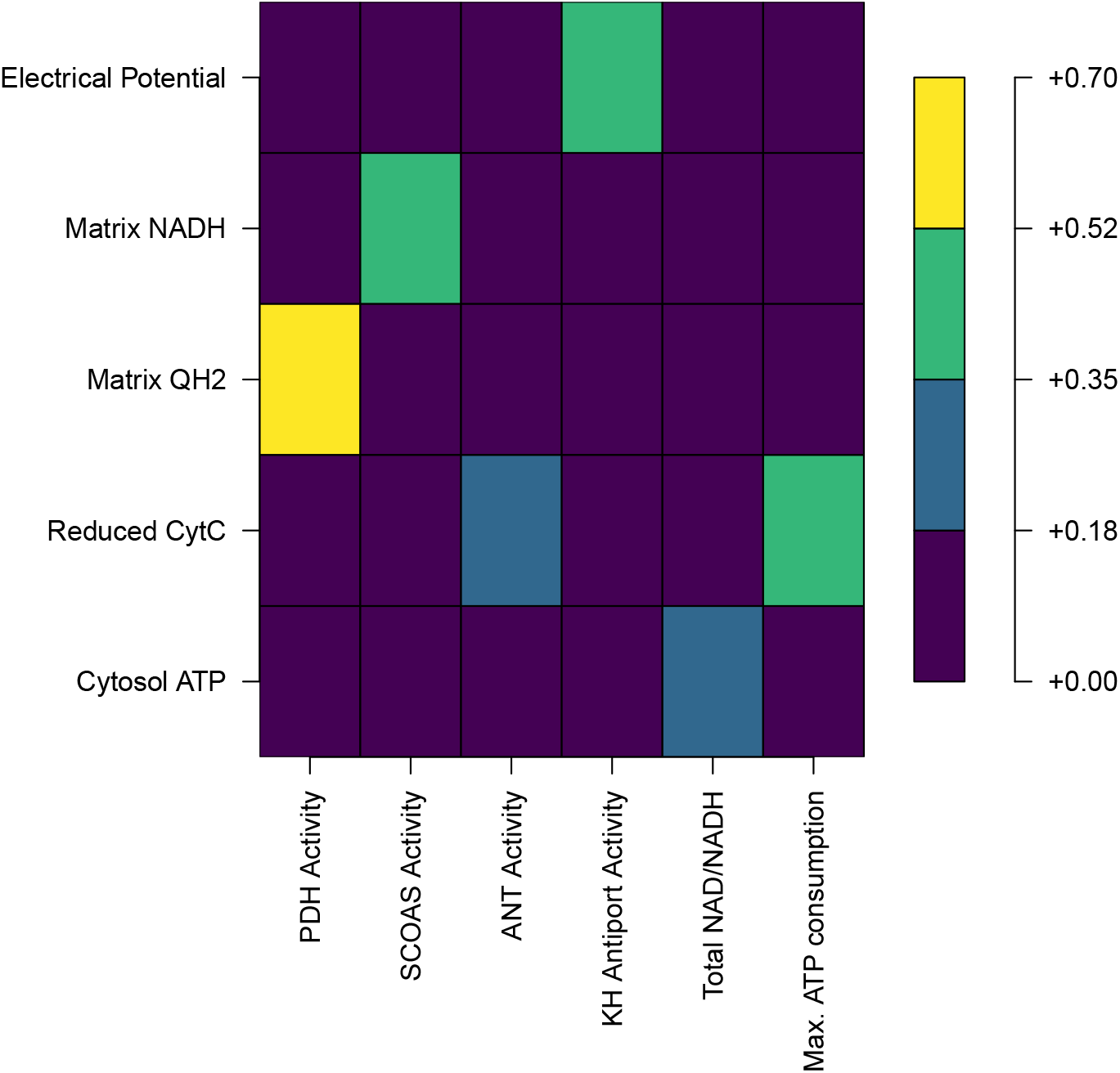
The sensitivity of important state variables to the most important parameters as calculated according to the Sobol method, not including interactions. CytC stands for cytochrome C, reduced cytochrome C refers to cytochrome C that has been donated an electron by Complex III.

**Figure 7:**
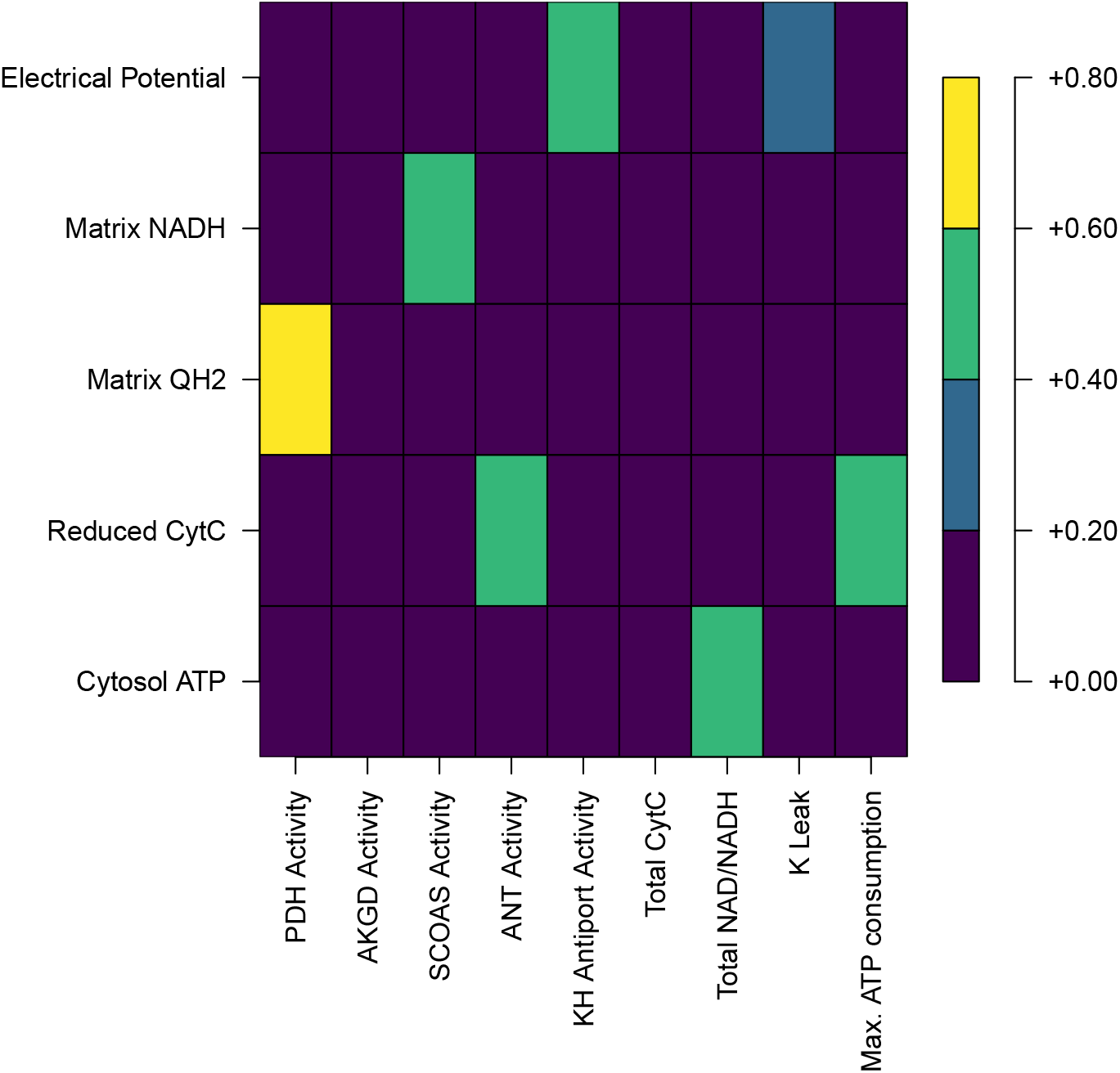
The sensitivity of important state variables to the most important parameters as calculated according to the Sobol method, taking into account two-way interactions. CytC stands for cytochrome C, reduced cytochrome C refers to cytochrome C that has been donated an electron by Complex III.

## References

[1] Beard Daniel A and Qian Hong. Chemical biophysics: quantitative analysis of cellular systems, volume 126. Cambridge University Press Cambridge, 2008.

[2] Chen Ying, Fry Brendan C, and Layton Anita T. Modeling glucose metabolism in the kidney. Bulletin of mathematical biology 78(6): 1318–1336, 2016.

[3] Cox Michael M and Nelson David L. Lehninger principles of biochemistry, volume 5. Wh Freeman New York, 2008.

[4] Edwards Aurélie, Castrop Hayo, Laghmani Kamel, Vallon Volker, and Layton Anita T. Effects of nkcc2 isoform regulation on nacl transport in thick ascending limb and macula densa: a modeling study. American Journal of Physiology-Renal Physiology 307(2): F137–F146, 2014.

[5] Edwards Aurélie, Palm Fredrik, and Layton Anita T. A model of mitochondrial o2 consumption and atp generation in rat proximal tubule cells. American Journal of Physiology-Renal Physiology 318(1): F248–F259, 2020.

[6] Fry Brendan C, Edwards Aurélie, and Layton Anita T. Impacts of nitric oxide and superoxide on renal medullary oxygen transport and urine concentration. American Journal of Physiology-Renal Physiology 308(9): F967–F980, 2015.

[7] Hoffman David L, Salter Jason D, and Brookes Paul S. Response of mitochondrial reactive oxygen species generation to steady-state oxygen tension: implications for hypoxic cell signaling. American Journal of Physiology-Heart and Circulatory Physiology 292(1): H101–H108, 2007.

[8] Jonker Ard, Geerts Willie JC, Charles Rob, Lamers Wouter H, and Van Noorden Cornelis JF. The dynamics of local kinetic parameters of glutamate dehydrogenase in rat liver. Histochemistry and cell biology 106(4): 437–443, 1996.

[9] Kohn Michael C and Garfinkel David. Computer simulation of metabolism in palmitate-perfused rat heart. ii. behavior of complete model. Annals of biomedical engineering 11(6): 511–531, 1983.

[10] Wu Fan, Yang Feng, Vinnakota Kalyan C, and Beard Daniel A. Computer modeling of mitochondrial tricarboxylic acid cycle, oxidative phosphorylation, metabolite transport, and electrophysiology. Journal of Biological Chemistry 282(34): 24525–24537, 2007.

